# Subsets of extraocular motoneurons produce kinematically distinct saccades during hunting and exploration

**DOI:** 10.1101/2024.08.12.607184

**Authors:** Charles K. Dowell, Thomas Hawkins, Isaac H. Bianco

## Abstract

Animals construct diverse behavioural repertoires by moving a limited number of body parts with varied kinematics and patterns of coordination. There is evidence that distinct movements can be generated by changes in activity dynamics within a common pool of motoneurons, or by selectively engaging specific subsets of motoneurons in a task-dependent manner. However, in most cases we have an incomplete understanding of the patterns of motoneuron activity that generate distinct actions and how upstream premotor circuits select and assemble such motor programmes. In this study, we used two closely related but kinematically distinct types of saccadic eye movement in larval zebrafish as a model to examine circuit control of movement diversity. In contrast to the prevailing view of a final common pathway, we found that in oculomotor nucleus, distinct subsets of motoneurons were engaged for each saccade type. This type-specific recruitment was topographically organised and aligned with ultrastructural differences in motoneuron morphology and afferent synaptic innervation. Medially located motoneurons were active for both saccade types and circuit tracing revealed a type-agnostic premotor pathway that appears to control their recruitment. By contrast, a laterally located subset of motoneurons was specifically active for hunting-associated saccades and received premotor input from pretectal hunting command neurons. Our data support a model in which generalist and action-specific premotor pathways engage distinct subsets of motoneurons to elicit varied movements of the same body part that subserve distinct behavioural functions.

## Introduction

Animals can move their individual body parts with varied kinematics and patterns of coordination to compose a broad variety of behaviours. This necessitates that different force profiles be generated by muscles, which in turn requires distinct patterns of motoneuron activity. Principles of motor control include size-ordered recruitment of motoneurons to produce increasingly forceful movements^1^, force trajectories encoded by dynamic population activity^2^, and task-specific recruitment^3^. However, in most instances we have an incomplete understanding of how movement diversity relates to population activity within motor pools and the circuit mechanisms by which premotor commands that encode specific kinematic variables or motor subroutines^4–7^ sculpt appropriate patterns of motoneuron activity.

The oculomotor system presents several advantages for elucidating how neural circuits generate a diverse motor repertoire. Several types of eye movement, with velocities spanning at least two orders of magnitude, support a variety of visual functions^8^ and are produced by only six eye muscles that are innervated by circumscribed pools of motoneurons in the brainstem. Recordings from these extraocular motoneurons has given credence to the idea of a ‘final common pathway’, in which different eye movement subsystems converge on a common population of motoneurons that participate in all types of eye movement^9^. However, this notion seems at odds with other anatomical and physiological evidence. Extraocular muscles are composed of diverse muscle fibre types, which are in turn innervated by motoneurons that vary in morphology, neurochemistry, physiological properties and afferent inputs^10^. Furthermore, physiological data has shown that the firing properties of single motoneurons can change in the context of distinct movement types ^11,12^ and that motoneurons can be divided into subgroups with distinct dynamic properties^13^. Such evidence suggests that motoneurons might show at least some degree of selective activity and/or recruitment to produce eye movements with distinct kinematics or subserving different visuomotor functions.

To try to resolve this conundrum, we took advantage of two closely related but kinematically and ethologically distinct types of saccadic eye movement, which are expressed by larval zebrafish. Saccades are brief but extremely rapid eye movements that enable animals to swiftly redirect gaze and are generated by stereotypical ‘phasic-tonic’ activity in extraocular motoneurons^14,15^. Although it has been assumed that all horizontal saccades are controlled by a common neural pathway^16^, we recently discovered that larval zebrafish generate different types of saccadic eye movement that are used in different behavioural contexts and which follow distinct kinematic rules^17^ (and [Fig.1]). Specifically, conjugate saccades, in which both eyes rotate in the same direction, are used to redirect gaze during exploration and recentre the eye during the optokinetic reflex. By contrast, convergent saccades, in which both eyes rotate nasally, play a specialised role during hunting to foveate prey targets and increase the extent and proximity of the binocular visual field ^17–20^. We found that adducting (nasally directed) horizontal saccades follow distinct velocity profiles and observe distinct relationships between eye velocity and saccade amplitude when they occur in the binocular context of a conjugate saccade versus a convergent saccade^17^. These kinematic differences indicate that these two types of adducting saccade are generated by different patterns of extraocular motoneuron population activity^21,22^.

**Figure 1:**
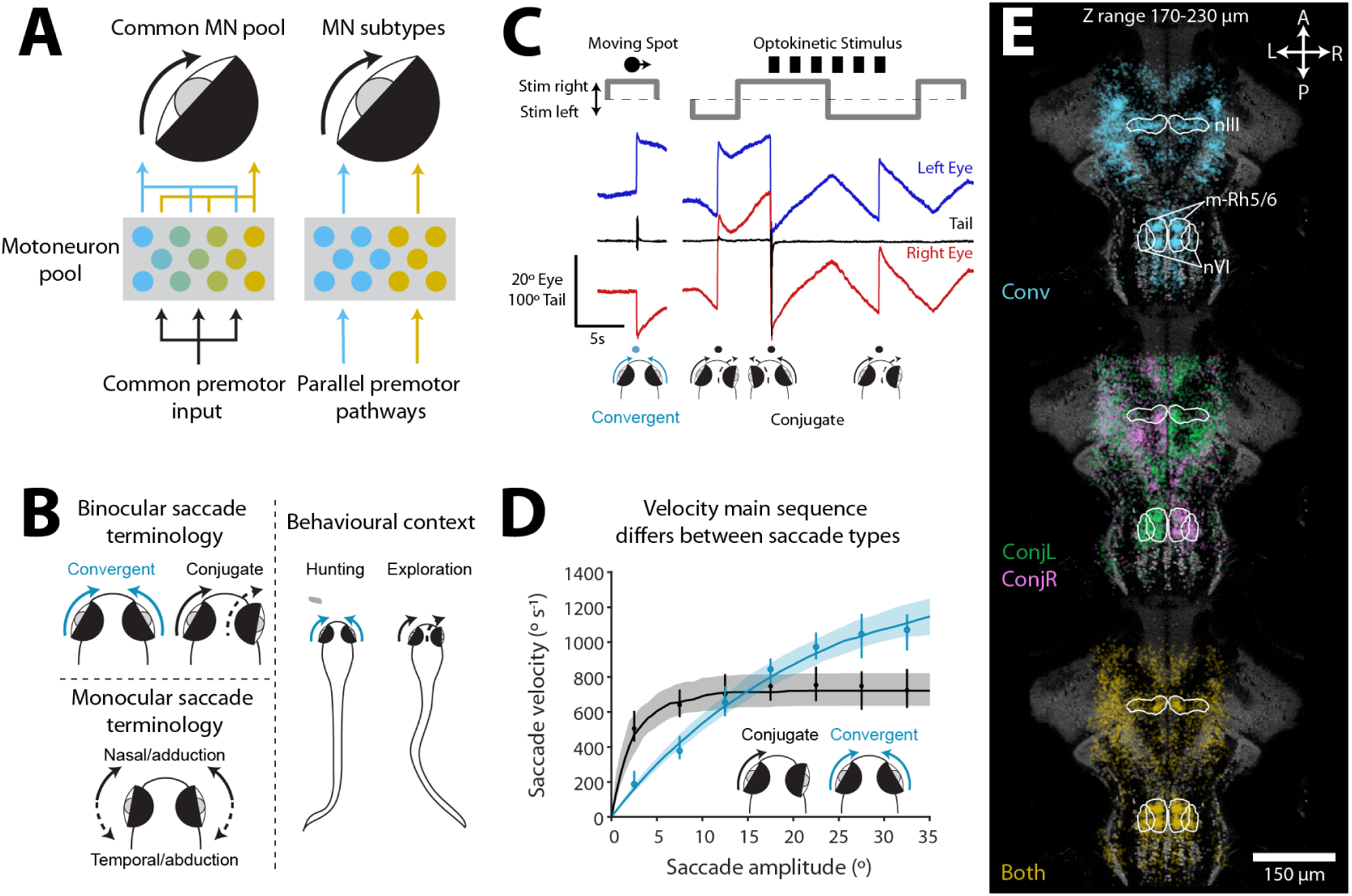
Saccade kinematics and oculomotor-tuned neurons. (A) Schematic illustrating two models for neural control of distinct movements of the same plant (in this case the eye). *Left:* Differences in spatio-temporal activity across a common population of motoneurons generates distinct muscle force profiles and motion kinematics. *Right:* Task-specific subsets of motoneurons generate distinct actions. Note that models are not mutually exclusive. (B) Key to saccade terminology. (C) Example behavioural tracking data from tethered animals undergoing 2-photon calcium imaging. Presentation of a prey-like moving spot evokes a hunting-related convergent saccade and a drifting grating evokes conjugate saccades (OKR fast phases). (D) Velocity main sequence for convergent and conjugate adducting saccades. Line and shading shows median *±* IQR across exponential fits for *N* = 96 eyes. For reference, median (*±* IQR) velocity for each amplitude bin is shown for data pooled across animals. (E) Oculomotor-tuned ROIs active for convergent (*Conv*) or leftwards/rightwards conjugate (*ConjL/R*) or both (*Both*) saccade types. Images show a single focal plane in the mid/hindbrain. For full data see [Fig.S2].

In this study, we leveraged these two saccade types to investigate two preeminent models for motor control: Namely, whether distinct but related movements are generated by a shared pool of motoneurons or by action-specific subsets of motoneurons and how such activation patterns might be programmed by premotor pathways [Fig.1A]. Using cellular-resolution calcium imaging, we discovered topographically organised and saccade type-specific activation of medial rectus motoneurons: Medially located cells were active during both saccade types whereas a laterally located subset were specifically engaged during convergent saccades. Electron microscopy revealed that this functional arrangement aligned with three subtypes of medial rectus motoneuron that differed in morphology and connectivity and suggested a synaptic mechanism for saccade type-specific motoneuron activation. Specifically, medially located motoneurons appeared to obtain the majority of their synaptic input via a remarkable ‘giant synapse’ from abducens internuclear neurons (INNs) and laser-ablations confirmed that INNs were required for both types of adducting saccade. By contrast, laterally located motoneurons received a differ-ent compliment of synaptic inputs, including monosynaptic innervation from pretectal hunting command neurons, thus identifying a circuit motif that links sensorimotor decision making to oculomotor output. In sum, our study supports a model in which parallel premotor pathways control the task-specific recruitment of subsets of motoneurons to generate kinematically distinct movements that subserve distinct ethological functions.

## Results

### Conjugate and convergent adducting saccades follow distinct kinematic rules and associate with different patterns of brainstem activity

Here, we used the fact that zebrafish produce kinematically distinct saccades when hunting versus during routine exploration to investigate how population activity across motoneurons generates distinct but related actions of the same body part [Fig.1A-B]. To examine neuronal activity during naturalistic behaviour, we combined 2-photon calcium imaging with high-speed recording of eye position and tail posture [Fig.1C]. During imaging, animals were shown prey-like moving spots to evoke hunting-related convergent saccades. In addition, drifting gratings were used to evoke the optokinetic response (OKR), which comprises ‘slow phase’ eye rotations in the direction of whole-field motion and intermittent conjugate saccades (‘fast phases’), which reset eye position when the eyes become eccentric in the orbit. We note that these fast phases display the same kinematic properties as other conjugate saccades, including those that occur spontaneously or in conjunction with swims ^17^.

Eye tracking data from tethered animals confirmed that adducting saccades displayed distinct kinematics when they occurred in the binocular context of a convergent saccade as compared to a conjugate saccade^17^. We define convergent saccades as binocular events during which both eyes make an adducting (nasally directed) saccadic movement and conjugate saccades as events where both eyes rotate in the same direction (thus one eye adducts and the other abducts), but not necessarily with equal amplitudes [Fig.1B]. Although the distribution of amplitudes and peak velocities was similar for convergent and conjugate adducting saccades [Fig.S1A], when we examined the velocity ‘main sequence’, which is a characteristic feature of saccadic eye movements and relates peak eye velocity to saccade amplitude^23^, we observed distinct main sequence relationships [Fig.1D]. At small amplitudes, conjugate adducting saccades were faster, but velocity saturated at *∼* 700*^◦^*/s at amplitudes exceeding 10*^◦^*. By contrast, convergent adducting saccades showed lesser saturation and eye velocity continued to scale with amplitude. These observations were supported by statistically distinct exponential fits^24,25^ describing the main sequence relationships (AIC = 97.7%, *p* = 7.5 *×* 10*^−^*^15^, *N* = 86 eyes, signed rank test versus single model). The eye also reached more nasal post-saccadic positions during convergent saccades (convergent = 11.1 *±* 0.2*^◦^*, conjugate = 5.5 *±* 0.2*^◦^*, mean *±* SEM across *N* = 152 eyes, *p* = 3.2 *×* 10*^−^*^50^, t-test; [Fig.S1A]). These observations and our previous characterisation^17^ suggest that distinct patterns of motoneuron activity are responsible for controlling adducting saccades in these two behavioural contexts.

To identify neural activity associated with the production of convergent and/or conjugate saccades, we performed calcium imaging in Tg(*elavl3*:H2B-GCaMP6s) transgenic animals, in which a genetically encoded calcium indicator is expressed broadly across the brain. In total we recorded the activity of 1, 124, 129 automatically segmented regions-of-interest (ROIs; corresponding to individual neurons) across the midbrain and hindbrain of 76 animals and then developed a two-stage analysis pipeline to identify ROIs that are putatively involved in generating saccadic eye movements [Fig.S1; Methods]. In this way, we identified 44, 332 ‘oculomotor-tuned’ ROIs (4.0 *±* 0.2% of the total 1.1M) and categorised them as active for either convergent (*Conv*) or conjugate (*Conj*) or both (*Both*) types of saccade. Brain volumes were registered to a reference brain (ZBB) ^26^ allowing us to examine the anatomical distribution of oculomotor-tuned ROIs in a standard coordinate space [Fig.1E, Fig.S2]. Of particular relevance to this study, oculomotor-tuned cells were enriched in the oculomotor (nIII) and abducens (nVI) nuclei, a region of medial rhombomere-5/6 (m-Rh5/6) that has previously been shown to be active during ipsiversive conjugate saccades^27–29^, and the pretectum adjacent to retinal arborization field 7 (AF7-Pt), which contains hunting command neurons^30^.

### Extraocular motoneurons are differentially recruited across saccade types

Saccades are generated by phasic-tonic activity in extraocular motoneurons, where a phasic ‘pulse’ of spiking rapidly accelerates the eye and firing rate then declines (‘glide’) to a tonic level (‘step’) to hold the eye in its new position^14^. The distinct velocity main sequence profiles indicate that the pulse component, which controls eye velocity, saturates for conjugate adducting saccades but continues to scale with amplitude during convergent saccades. This might be due to differences in net population activity within a common pool of motoneurons, or might arise from recruitment of different subsets of motoneurons between saccade types. We therefore examined activity in the oculomotor (nIII) and abducens (nVI) nuclei, which contain medial rectus motoneurons (MRMNs), lateral rectus motoneurons (LRMNs) and abducens internuclear neurons (INNs) that have well established roles in generating horizontal eye movements [Fig.2A].

**Figure 2:**
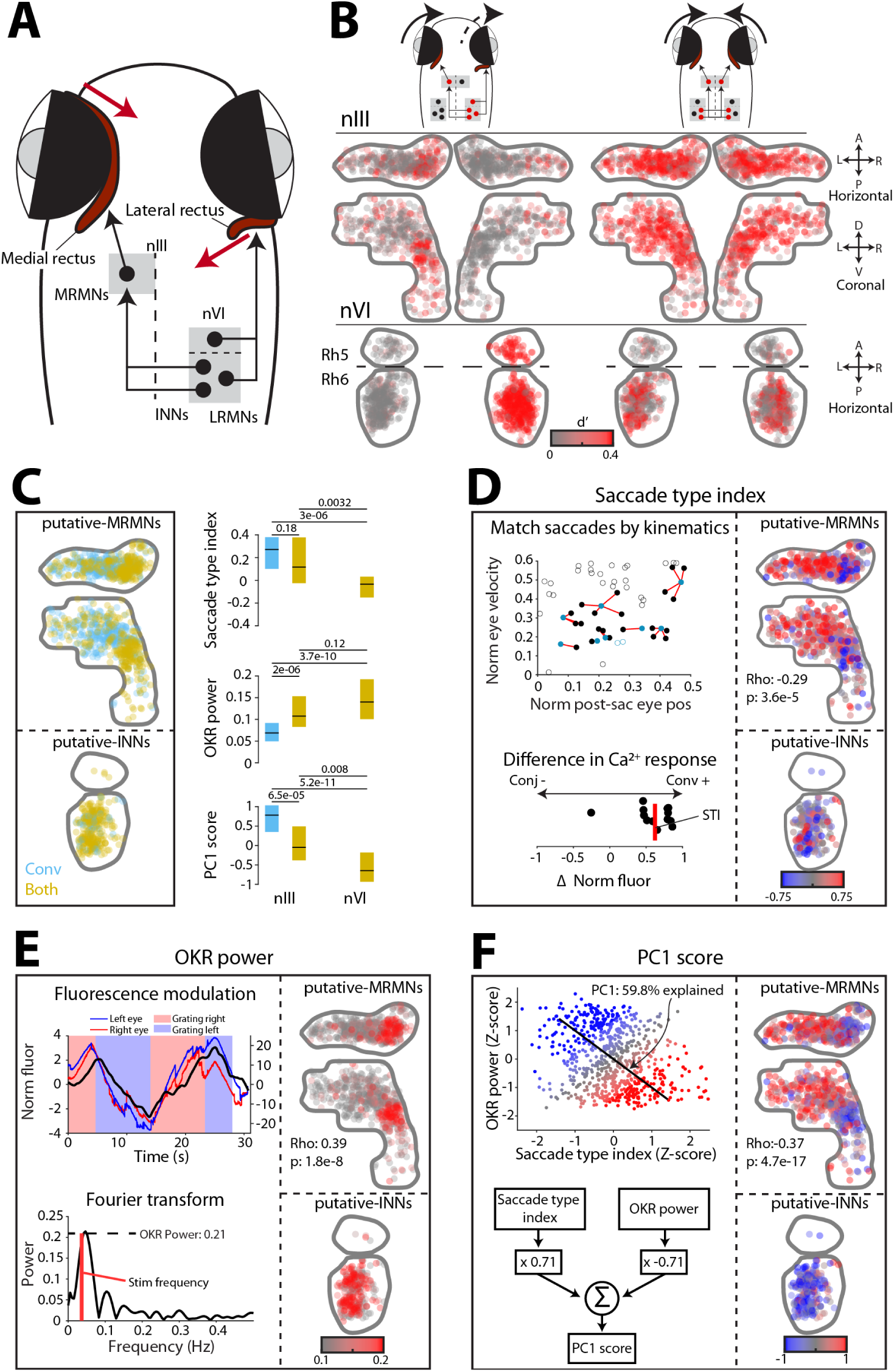
Subsets of extraocular motoneurons are engaged during conjugate versus convergent saccades. (A) Horizontal gaze control by neurons in oculomotor (nIII) and abducens (nVI) nuclei. (B) Oculomotor-tuned ROIs coloured by saccade-triggered activity (d’) for rightwards conjugate saccades and convergent saccades (1,720 ROIs from 68 fish). (C) Oculomotor-tuned ROIs in nIII (pMRMNs, *N* = 538) and nVI (pINNs, *N* = 292) classified by saccade-type activation. In this and subsequent panels, right-sided ROIs have been reflected onto the left. *Right:* Functional metrics (median with IQR) for pMRMNs and pINNs. *p*-values from Kruskal-Wallis with Dunn-Sidak post-hoc test. (D) Saccade type index. *Left:* Process of computing saccade type index for an example neuron. First, each convergent saccade (blue) is matched to a conjugate saccade (black) having similar peak velocity and post-saccadic eye position. Matched pairs indicated with red lines. Saccade type index is then the median difference in saccade-triggered fluorescence modulation across all matched pairs. *Right:* pMRMNs in nIII and pINNs in nVI, colour-coded by saccade type index (*N* = 743 ROIs). (E) OKR power. *Left:* Process of computing OKR power, illustrated for an example cell. First, median fluorescence modulation is computed during optokinetic stimulation. Then the power spectrum of the fluorescence signal is computed and OKR power is measured at the stimulus frequency (red line). *Right:* As per (D) for OKR power (*N* = 830 ROIs). (F) PC1 score. *Left:* Illustration of PC1. *Right:* As per (D) for PC1 score (*N* = 743 ROIs).

Oculomotor-tuned cells showed distinct patterns of saccade-triggered activity modulation (d’) during conjugate versus convergent saccades [Fig.2B]. For conjugate saccades, we observed the expected pattern of lateralised activity, which was consistent with unilateral activity in MRMNs (ipsilateral to the adducting eye) as well as LRMNs and INNs in the contralateral abducens (i.e. ipsilateral to the abducting eye). By contrast, convergent saccades were associated with symmetric activity, consistent with bilateral activation of INNs driving bilateral recruitment of MRMNs to produce adduction of both eyes. The identities of MRMNs, INNs and LRMNs were supported by analysis of the eye–direction tuning of individual ROIs. By fitting rectilinear functions relating saccade-triggered fluorescence to post-saccadic eye position [Fig.S3A], we found that in abducens, *Both* ROIs were tuned to adduction of the contralateral eye, identifying these cells as putative INNs (pINNs), whereas *Conj* ROIs were activated for abduction of the ipsilateral eye, as expected for LRMNs [Fig.S3B-C]. In nIII, *Both* and *Conv* ROIs were tuned to adduction of the ipsilateral eye, as expected for MRMNs^31^ and are thus designated putative MRMNs (pMRMNs). Moreover, these cells were located in the dorsal subdivision of nIII, where retrograde tracing from eye muscles has localised MRMNs in larval zebrafish^32^. *Conj* ROIs in nIII occupied scattered locations and showed no systematic nasal/temporal preference [Fig.S3B] and are not considered further.

In support of the hypothesis that recruitment of distinct subsets of motoneurons might underlie distinct saccade kinematics, we observed strikingly different patterns of pMRMN activity during conjugate versus convergent saccades. Specifically, while conjugate saccades were associated with activity that was restricted to a relatively compact dorso-medial region, convergent saccade activity extended much more broadly across dorso-lateral nIII [Fig.2B]. Very similar patterns were observed when saccades were binned according to amplitude or post-saccadic eye position [Fig.S3D], supporting the idea that these differences in recruitment are not due to systematic differences in kinematics between our saccade samples, but are instead an explicit function of saccade *type*. This pattern could also be seen in maps of ROIs labelled by their saccade type recruitment [Fig.2C]: *Both* ROIs were restricted to dorso-medial nIII but *Conv* ROIs extended across a broader dorso-lateral region. In sum, these results are consistent with differential recruitment of MRMNs generating convergent versus conjugate adducting saccades.

### Functional topography within oculomotor nucleus

Next, we analysed the response properties of individual neurons, which provided further support for saccade type-specific activity and revealed a topographic organisation of functional properties across the pMRMN population.

To assess the extent to which a cell’s activity was specifically modulated as a function of saccade type, independent of its eye position or velocity sensitivity, we developed a metric we termed ‘saccade type index’. Specifically, we computed the median difference in saccade-triggered calcium response between conjugate versus convergent saccades that were pairwise matched for similar post-saccadic eye positions and peak velocities [Fig.2D]. A positive index indicates a greater calcium response for convergent saccades, whereas a negative index indicates a greater response during conjugate saccades. Maps of pMRMNs coded by saccade type index revealed a clear topography within nIII [Fig.2D]. In dorso-medial nIII the mean index was close to zero, indicating no overall preference for either saccade type. However, in dorso-lateral nIII the majority of cells had positive index, indicating greater modulation during convergent saccades. This pattern mirrors the anatomical segregation of *Both* and *Conv* ROIs [Fig.2C] and supports the notion that MRMNs are recruited in a saccade type-specific manner and organised with functional topography within the oculomotor nucleus.

Functional topography was further evidenced by an independent functional metric, ‘OKR power’, which quantified the modulation of single-neuron fluorescence during slow phase eye movements [Fig.2E; Methods]. To provide a compact summary of single-cell functional properties, we used principal component analysis to compute a weighted sum of saccade type index and OKR power [Fig.2F]. The resulting ‘PC1 score’ increased significantly across the medial→lateral axis of nIII and consequently, laterally located *Conv* pMRMNs had significantly higher scores than medially localised *Both* pMRMNs [Fig.2C,F]. In abducens, *Both* ROIs (putative INNs, see above) had low PC1 scores that were most similar to *Both* pMRMNs, suggesting INNs might comprise the dominant afferent input to MRMNs in dorso-medial nIII.

In summary, our data support a model in which INNs recruit functionally similar MRMNs in dorso-medial nIII during both types of adducting saccade but during convergent saccades, additional MRMNs are recruited in dorso-lateral nIII.

### Three subtypes of MRMN with distinct morphology and afferent connectivity

We next asked if there were structural correlates of these functional differences by examining neuronal morphology and synaptic connectivity.

We started by examining INNs, as these neurons make excitatory, monosynaptic connections onto MRMNs^33–37^ and our imaging data indicate are active for both types of saccade. First, we visualised INN projections by performing 2-photon photoactivation of paGFP in abducens of Tg(*Cau.Tuba1*:c3paGFP)a7437; Tg(*elavl3*:jRCaMP1a)jf16 double transgenic animals [Fig.3A]. As expected^38,39^, axons of photolabelled INNs crossed the ventral midline at the level of nVI and ascended in the contralateral medial longitudinal fasciculus before arborising in nIII [Fig.3B]. We noticed that INN axonal boutons in the oculomotor nucleus varied in size [Fig.3B], with particularly large boutons (*≥* 4 *µm*^2^ cross-sectional area) occupying a medial region that corresponded to the location of *Both* pMRMNs [Fig.3C], whilst smaller boutons were distributed over a broader medio-lateral extent.

**Figure 3:**
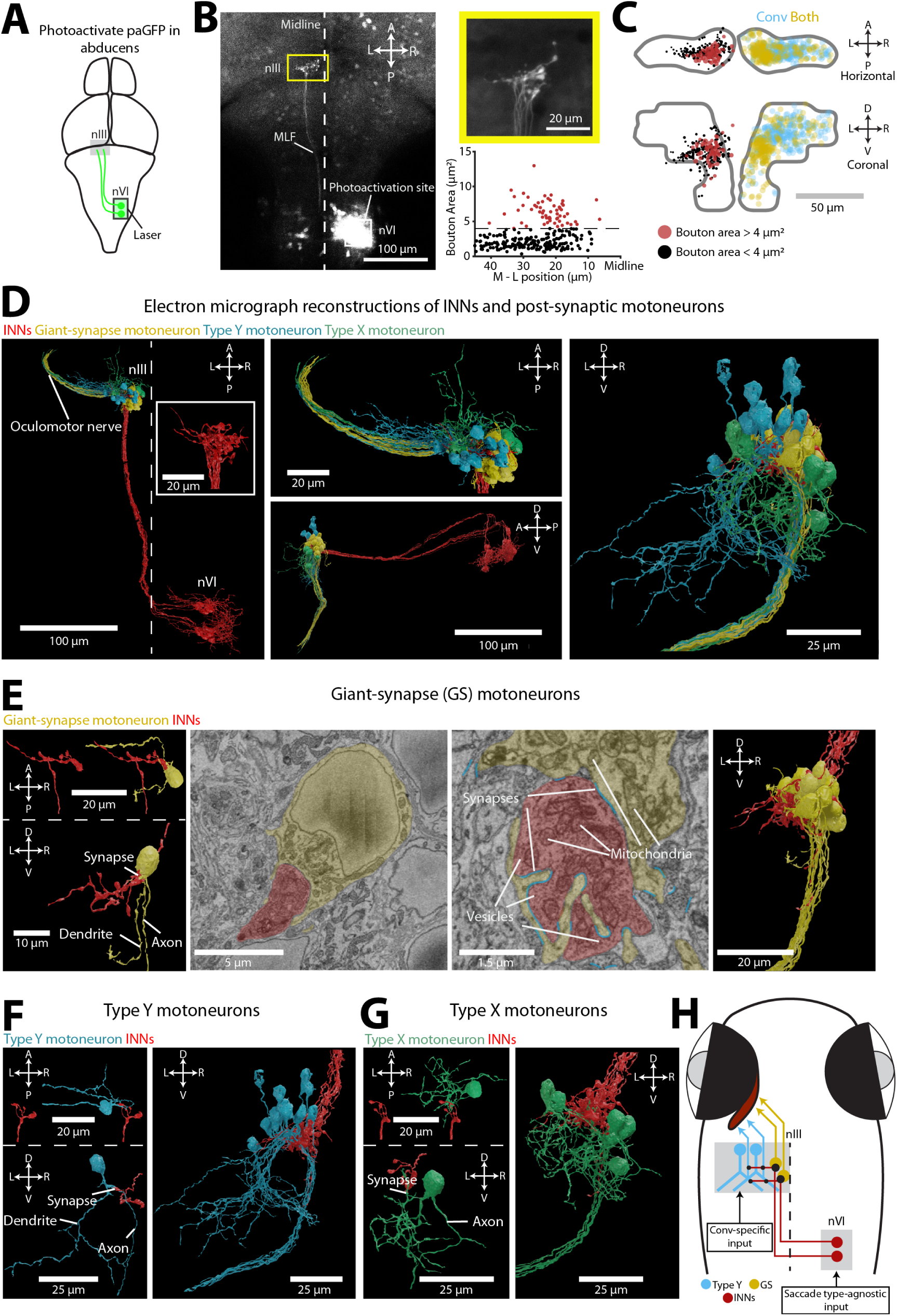
Three subtypes of medial rectus motoneuron. (A) Schematic of paGFP photoactivation experiment with 40 *×* 40 *×* 20 *−* 30 *µ*m target volume in nVI. (B) *Left:* Example paGFP labelling of INN projections from abducens to oculomotor nucleus. *Right, top:* High magnification image of INN axon terminals. *Right, bottom:* Cross-sectional area of axonal boutons versus mediolateral position in nIII (269 boutons from 7 animals). (C) Bouton locations in left nIII and oculomotor-tuned ROIs (shown on right). Circles indicate bouton locations and are scaled according to cross-sectional area. (D) Ultrastructural reconstructions of 15 INNs along with 35 post-synaptic motoneurons. Midline indicated by dashed line. (E) Giant-synapse motoneurons. *Left:* 3D renderings of an INN terminal arbor and post-synaptic giant-synapse motoneuron. *Middle:* Electron micrographs of the axon terminal (red) and post-synaptic motoneuron (yellow). Automatically detected synapses shown by blue lines. *Right:* 3D rendering of all 15 giant-synapse motoneuron somata, each associated with one INN. (F–G) 3D reconstructions of single (left) and all (right) Type Y (F, 12 cells) and Type X (G, 6 cells) motoneurons. (H) Circuit model (see text for explanation).

This observation led us to hypothesise that distinct patterns of synaptic connectivity between INNs and MRMNs might accompany the topographic gradient of motoneuron functional properties. To investigate, we used a publicly available whole-brain serial-blockface scanning electron micrography dataset^40^ to trace 15 INNs in one hemisphere, along with 35 post-synaptic neurons. All of the post-synaptic neurons extended an axon in the third cranial nerve and because they also receive synaptic input from INNs they are assumed to be MRMNs. Ultrastructural reconstructions revealed that individual INNs had multiple synaptic boutons within nIII. Remarkably, every INN established a ‘giant synapse’ with a single MRMN (*N* = 15) [Fig.3E, Fig.S4A]. This synapse was formed between one especially large INN bouton and claw-like invaginations of the soma of the post-synaptic motoneuron and to our knowledge has not previously been described in any species. To better resolve the synaptic contacts, we obtained additional transmission electron micrographs in nIII, which revealed multiple post-synaptic densities at the apposition between large axon terminals and the soma of putative motoneurons [Fig.S5], indicating that these ‘giant synapses’ contain multiple sites of neurotransmission. The post-synaptic MRMNs (which we refer to as giant-synapse (GS) motoneurons) otherwise had small and simple dendritic trees [Fig.3E, Fig.S4A], suggesting that the majority of their synaptic input derives from the giant synapse in a one-to-one connectivity motif with an INN. We also reconstructed two other motoneurons with simple dendritic arbors that formed large claw-like post-synaptic contacts with multiple INN boutons and had somata that sat adjacent to the other giant-synapse MRMNs [Fig.S4B].

In addition to the giant synaptic terminal, all INNs additionally had smaller synaptic boutons that contacted MRMNs that we classified into two types based on morphology: Type X and Type Y. Type Y motoneurons had large, ventrally directed dendritic arbors with prominent Y-shaped branches (*N* = 12 [Fig.3F, Fig.S4C], plus *N* = 5 cells in the opposite brain hemisphere [Fig.S4E]). INNs did not synapse onto these distal dendrites but rather onto the axons or proximal dendrites of the cells [Fig.3F, Fig.S4C]. Type X motoneurons (*N* = 6) were characterised by dendritic arbors that occupied medial and anterior portions of nIII and INNs synapsed onto their somata or distal dendrites [Fig.3G, Fig.S4D].

The cell bodies of giant-synapse, Type Y and Type X MRMNs occupied distinct positions within the oculomotor nucleus [Fig.3D] and the distribution of giant-synapse and Type Y motoneurons in particular bore striking resemblance to the medio-lateral organisation of functionally identified pMRMNs [compare Fig.3C and D]. Giant-synapse motoneurons occupied dorso-medial locations, similar to *Both* pMRMNs that were active for conjugate and convergent saccades and have PC1 scores more similar to INNs. By contrast, Type Y motoneurons were located dorso-laterally, consistent with pMRMNs that had high saccade type indices and were preferentially recruited during convergent saccades.

Together, these observations support a model in which topographically organised synaptic connectivity contributes to distinct functional properties and saccade type-specific recruitment of MRMNs [Fig.3H]. In this model, the giant synapses between INNs and medially located GS motoneurons provide a strong feedforward relay of INN activity such that both cell types share similar functional profiles and are active for both types of saccade. Type Y motoneurons also receive excitatory input from INNs on their axons and proximal dendrites, but likely receive additional afferent input that underlies the convergent saccade-specific activity observed in lateral pMRMNs.

### Abducens internuclear neurons are necessary for both convergent and conjugate adducting saccades

Our model predicts that INNs provide a significant synaptic input to MRMNs for the production of both conjugate and convergent adducting saccades. To test the necessity for INN innervation, we performed two loss-of-function experiments designed to abrogate INN input.

First, we used a pulsed infrared laser to ablate the somata of functionally identified INNs [Fig.4A]. To do this, we first used 2-photon calcium imaging in abducens to identify neurons that were active during convergent saccades, reasoning that these cells should correspond to INNs (rather than LRMNs, see above). After ablation of functionally identified pINNs (14 *±* 3 cells in *N* = 7 animals), we observed a substantial impairment in post-saccadic position and velocity of the contralateral eye during both conjugate and convergent adducting saccades [Fig.4B-C]. The contralateral eye also obtained a more eccentric position following temporal saccades, likely as a result of reduced tone in MRMNs. Finally, we observed a small increase in nasal velocity of the ipsilateral eye, perhaps due to unintended damage to LRMNs [Fig.4C].

**Figure 4:**
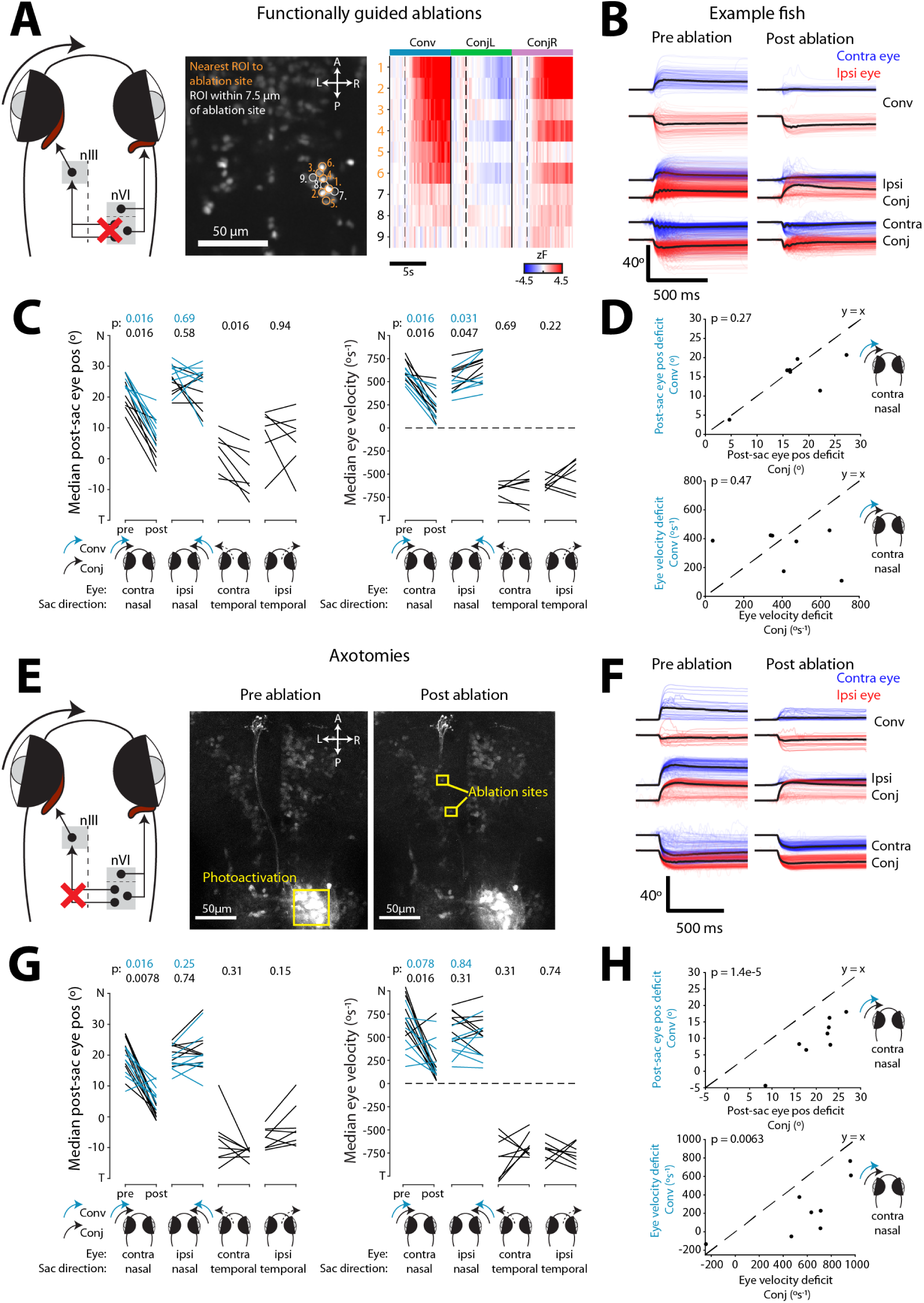
Abducens internuclear neurons are necessary for both conjugate and convergent adducting saccades. (A) *Left:* Schematic of ablations. *Right:* Example plane showing locations of functionally targeted ROIs and their median saccade-triggered activity. Dashed line indicates saccade time. (B) Eye position traces for saccades pre-and post-ablation, for an example fish. Black lines show median across trials. *Ipsi* and *Contra* refer to eye (ipsilateral or contralateral) and saccade direction (ipsiversive or contraversive), with respect to the ablation site. (C) Post-saccadic eye position and eye velocity for conjugate and convergent saccades, pre-and post-ablation. Median across saccades for each of *N* = 7 animals. (D) Deficits in post-saccadic eye position (top) and velocity (bottom) for conjugate versus convergent nasal saccades of the contralateral eye. Deficits computed as differences between medians pre-and post-ablation. (E) *Left:* Schematic of axotomies. *Right:* Example axotomy of photolabelled INN axons in MLF. (F–H) As per B–D, for axotomies (*N* = 8 animals). *p*-values in C, G are for signed-rank tests and in D, H for t-tests.

Second, we performed laser axotomies of INN projections in the medial longitudinal fasciculus, which we targeted following photolabelling with paGFP (*N* = 8 animals) [Fig4.E]. Axotomy caused similar deficits in adducting saccades of the contralateral eye [Fig.4F-G], although the deficit was weaker for convergent saccades [Fig.4H]. In part, we suspect this is due to the ablations failing to cut all INN axons and the stronger effect on conjugate adducting saccades may indicate a greater dependence of this saccade type on INN innervation.

In summary, loss-of-function experiments support a model in which INN input to MRMNs is required for both convergent and conjugate adducting saccades with normal kinematics. The lesser impact of INN axotomies on convergent saccades is in line with our hypothesis that additional afferent input is involved in generation of this saccade type.

### Medial rhombomere-5/6 makes functionally distinct connections to INNs and Type Y motoneurons

We next sought to identify premotor input to INNs, which our model predicts would be involved in generating both saccade types, as well as parallel inputs to Type Y motoneurons that should play a specific role in convergent adducting saccades. Horizontal saccades are triggered by disinhibition of burst neurons in the paramedian pontine reticular formation and medullary reticular formation, which in turn provide eye velocity signals to abducens^14^. Because saccadic burst neurons have been optogenetically mapped to rhombomere 5 in larval zebrafish^41^, we focussed on m-Rh5/6, where we observed a concentration of neurons that were active during convergent saccades (*Conv*) or both saccade types (*Both*) [Fig.1, Fig.S2].

Neurons located in the dorsal part of m-Rh5/6 provided direct synaptic input to ipsilateral INNs. We showed this by first photoactivating paGFP in abducens and subsequently identifying retrogradely labelled cell bodies [Fig.5A,B]. Photolabelled somata were observed in various mid/hindbrain regions, including a high density of cells in the dorsal region of m-Rh5/6, ipsilateral to the photoactivation site [Fig.5C]. To verify this putative connection, we reconstructed cells from the Svara et al, 2022^40^ EM dataset. By tracing cells from their presynaptic terminals with INNs back towards their somata, we identified 5 neurons in dorsal m-Rh5/6 [Fig.5D], thus confirming that neurons in this region make monosynaptic connections onto INNs.

**Figure 5:**
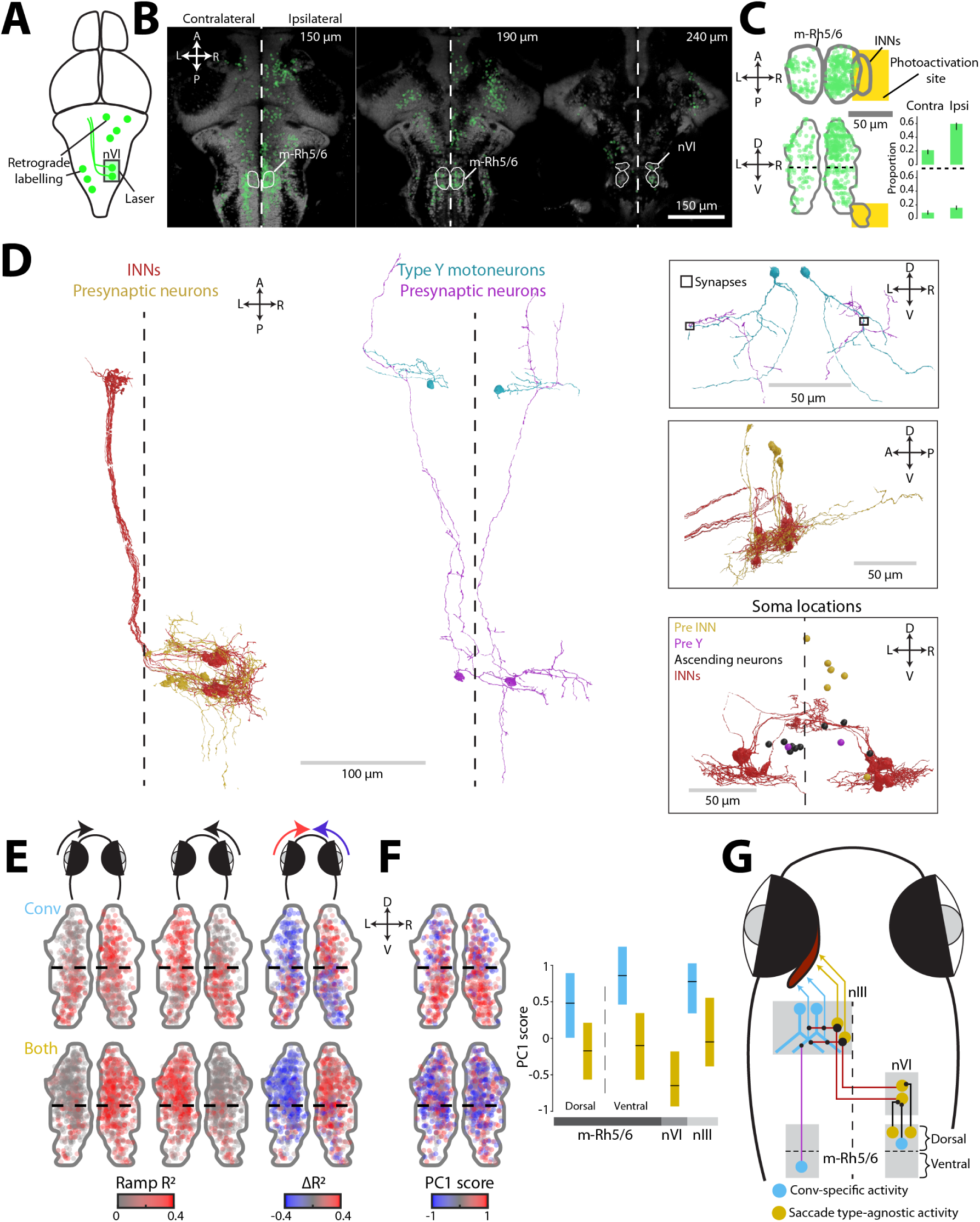
Medial rhombomere 5/6 makes functionally distinct connections to INNs and Type Y motoneurons. (A) Schematic of retrograde photolabelling from abducens. (B) Retrogradely labelled somata plotted on horizontal planes in ZBB reference space (4,063 neurons from 7 animals). (C) Locations of retrogradely labelled neurons in m-Rh5/6 (440 neurons). Inset shows proportion of cells labelled in dorsal and ventral domains of m-Rh5/6, ipsi-and contralateral to photoactivation site (median (IQR), *N* = 7 animals). (D) Ultrastructural reconstructions of six neurons in m-Rh5/6 that are presynaptic to INNs (left) and two cells that are presynaptic to Type-Y motoneurons (right). Inset boxes show synapses, alternate views and soma locations of other ascending neurons from m-Rh5/6. (E) Topography in eye–direction tuning along dorsoventral axis of m-Rh5/6 (891 *Conv*, 1,447 *Both* ROIs from 74 fish). *Left, middle:* Rectilinear fit *R*^2^ for nasal rotation of the left and right eye. *Right:* Cell-wise difference in *R*^2^ for left versus right eye nasal rectilinear fits. There is a switch in tuning from contralateral to ipsilateral eye along the dorsal to ventral axis. (F) PC1 score for oculomotor-tuned ROIs in m-Rh5/6 (814 *Conv*, 1,299 *Both* ROIs from 71 fish). Plot shows median (IQR) PC1 score across ROIs in m-Rh5/6, as well as pINNs and pMRMNs (from Fig.2) for comparison. (G) Circuit model (see text for explanation).

Ultrastructural data also revealed that neurons in ventral m-Rh5/6 provide afferent input directly to Type Y motoneurons. We showed this by reconstructing cells with somata in ventral m-Rh5/6 and in so doing identified 11 neurons with ipsilateral ascending projections to the caudal midbrain [Fig.5D, Fig.S6A]. For two of these neurons, we confirmed synaptic connections onto the distal Y-shaped dendrites of Type Y motoneurons [Fig.5D], thereby identifying an INN-independent afferent input to this subtype of MRMN. We also identified 11 additional neurons, in diverse midbrain and hindbrain locations, that were presynaptic to Type Y motoneurons [Fig.S6B], indicating that they receive several sources of innervation.

Based on these findings that dorsal m-Rh5/6 innervates INNs and ventral m-Rh5/6 innervates Type Y motoneurons, we hypothesised that there should be a dorso-ventral topography of functional properties in m-Rh5/6. Indeed, when we fit rectilinear functions to oculomotor-tuned ROIs, we observed a switch in eye–direction tuning along the dorso-ventral axis [Fig.5E]: Neurons in dorsal m-Rh5/6 were tuned to adduction of the contralateral eye, consistent with ipsilateral input to INNs (which in turn innervate contralateral MRMNs), whereas *Conv* ROIs in ventral m-Rh5/6 were tuned to adduction of the ipsilateral eye, consistent with projections from this region to ipsilateral Type Y MRMNs. In addition, OKR power decreased and saccade type index increased from dorsal to ventral m-Rh5/6 [Fig.S6C]. As a result, *Conv* ROIs in ventral m-Rh5/6 had high PC1 scores and so appeared functionally similar to *Conv* pMRMNs in dorso-lateral nIII. By contrast, *Both* ROIs, especially in dorsal m-Rh5/6, had lower PC1 scores, similar to pINNs and *Both* pMRMNs in dorso-medial nIII [Fig.5F].

Together, these data support a model in which m-Rh5/6 contributes to seperate premotor channels to recruit the subsets of motoneurons that produce conjugate and convergent adducting saccades [Fig.5G]. Cells in dorsal m-Rh5/6 are putative saccadic burst neurons that innervate INNs to generate both types of saccade. By contrast, neurons in ventral m-Rh5/6 make direct connections onto Type Y motoneurons and appear to provide parallel, convergent saccade-specific, signals.

### Medial rhombomere-5/6 is required for adducting saccades and its activation drives nasal eye movement

We next performed gain-and loss-of-function experiments to probe the causal role for m-Rh5/6 neurons in the generation of saccadic eye movements.

First, we performed functionally guided laser-ablations [Fig.6A]. Removal of saccade-active m-Rh5/6 neurons resulted in eye velocity deficits during adducting saccades of the contralateral eye [Fig.6B], similar to ablations of INNs. Deficits in post-saccadic position scaled with the number of ablated neurons and were very similar for conjugate and convergent saccades [Fig.6C]. The effect of ablations on the ipsilateral eye was more complex, producing less robust velocity and position deficits [Fig.6B]. Moreover, the effect on post-saccadic eye position showed substantial variability between saccade types [Fig.6D]. To attempt to explain this variability, we analysed the functional properties of ablated neurons and observed that the difference in effect size between convergent versus conjugate saccades (‘Conv-Conj residual’) was linearly related to the median saccade type index of the ablated neurons [Fig.6D]. Together, these data support a model in which m-Rh5/6 input to ipsilateral INNs is required to drive both types of adducting saccade in the contralateral eye. Although nasal saccades of the ipsilateral eye were less impacted, the greater sensitivity of convergent saccades to loss of neurons with high saccade type index is in line with a convergent-specific ipsilateral input onto Type Y motoneurons.

**Figure 6:**
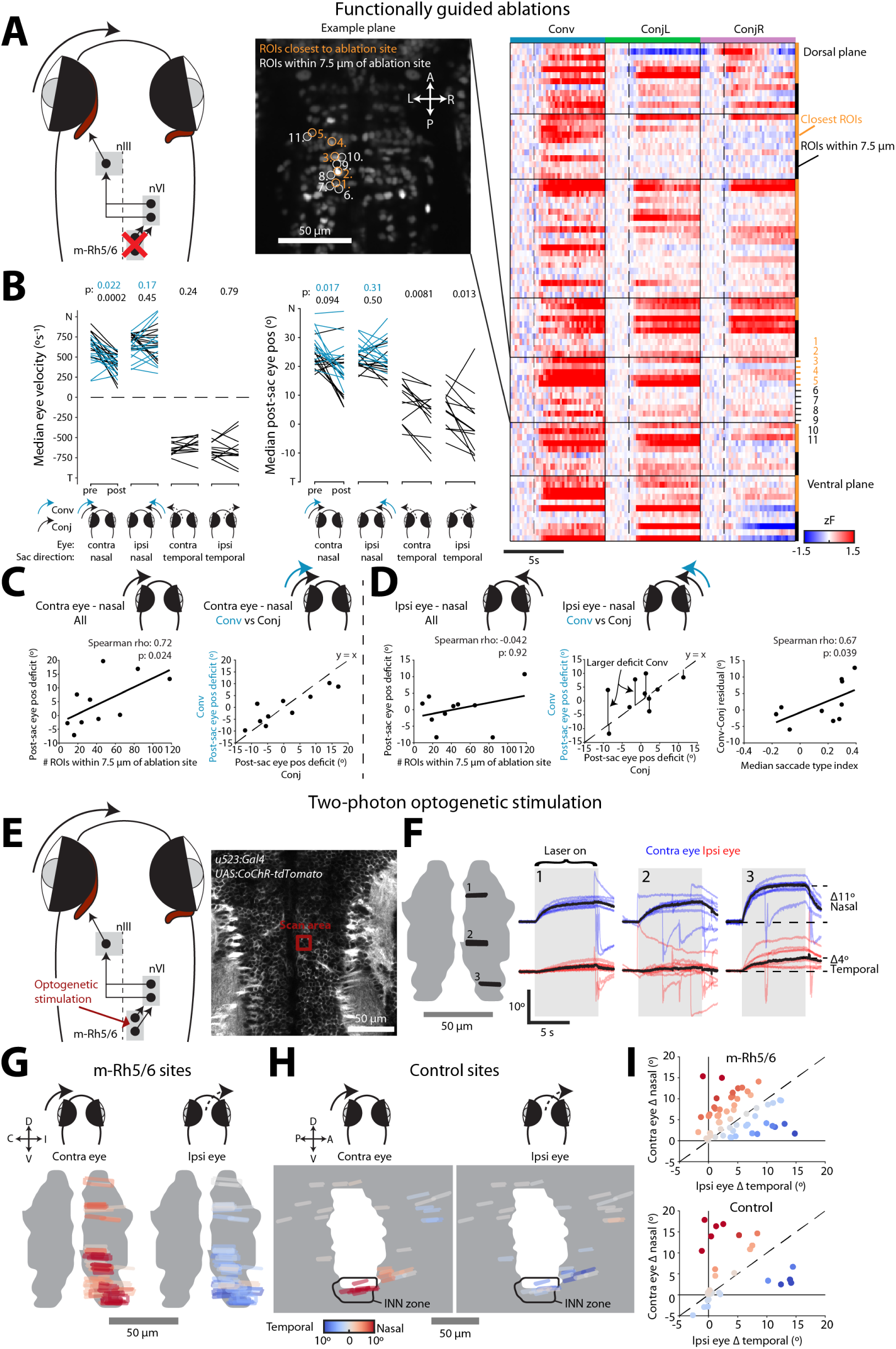
Ablation and optogenetic activation support a role for m-Rh5/6 in control of adducting saccades. (A) *Left:* Schematic of ablations. *Right:* Functionally targeted ROIs from one example focal plane and saccade-triggered activity for all targeted ROIs across seven focal planes, in the same animal. (B) Eye velocity and post-saccadic eye position pre-and post-ablation (medians across saccades for each of *N* = 13 animals, signed-rank test). (C) Post-saccadic eye position deficit for adducting saccades of the eye contralateral to ablation site. *Left:* Deficit versus number of ROIs within 7.5 *µ*m of ablation site. *Right:* Comparison of deficit for convergent versus conjugate saccades. (D) Post-saccadic eye position deficit for adducting saccades of the eye ipsilateral to ablation site. *Left, middle:* as per C. *Right:* Difference between deficit for convergent and conjugate saccades (Conv-Conj residual) versus median saccade type index of ablated cells. (E) Schematic of 2-photon optogenetic stimulation of m-Rh5/6 neurons and example scan site. (F) Eye position traces from an example animal for stimulation of three sites (indicated on a frontal view of m-Rh5/6 mask). Black lines show medians across trials. (G–H) Stimulation sites (G; *N* = 60 from 7 fish) and control sites (H; *N* = 36 from 8 fish), colour-coded by change in position of the contralateral (left panels) and ipsilateral (right panels) eye. All sites registered to ZBB reference space and depicted on the right hemisphere. (I) Pairwise comparison of change in position of the contra-versus ipsilateral eye. Points coloured according to distance from *y* = *x* line (dashed diagonal).

Next, we used multiphoton optogenetics to show that activating m-Rh5/6 neurons was sufficient to evoke nasal eye rotations. Specifically, we used Tg(*u523*:Gal4);Tg(UAS:CoChR-tdTomato) transgenic animals, in which the excitatory opsin CoChR is broadly expressed in mid/hindbrain and performed 2-photon photostimulation at localised sites within m-Rh5/6 [Fig.6E]. All sites evoked adduction of the contralateral eye, albeit to variable extents, compatible with m-Rh5/6 innervating INNs [Fig.6F-G]. By contrast, control stimulations in loci surrounding m-Rh5/6 produced little eye movement, except for stimulation in the region of INN somata which, as expected, evoked robust adduction of the contralateral eye [Fig.6H]. Most photostimulations in m-Rh5/6 also evoked abducting movements of the ipsilateral eye. Moreover, across different sites, we observed independent variation in the effects on the contralateral versus ipsilateral eye, suggesting a largely monocular organisation of oculomotor commands within m-Rh5/6 [Fig.6I].

In sum, anatomical, functional imaging, and gain-and loss-of-function experiments support the existence of two output pathways from m-Rh5/6. Dorsal m-Rh5/6 innervates INNs and this pathway appears both necessary and sufficient for adducting saccades of the contralateral eye. Ventral m-Rh5/6 contains neurons that directly innervate ipsilateral Type Y motoneurons. However, the weaker effects of ablation and optogenetic stimulation on adduction of the ipsilateral eye seem compatible with this pathway operating in parallel to INN input and being only one of several potential sources of convergent saccade-specific innervation of motoneurons.

### Pretectal command neurons make synaptic connections onto oculomotor tar-gets

Zebrafish use conjugate saccades to shift gaze during routine exploration of their environment and to recentre the eye during OKR, whereas convergent saccades are deployed during hunting to binocularly foveate prey^17^. How do descending commands interface with oculomotor circuits to generate appropriate saccade types in distinct ethological contexts?

We previously identified a population of neurons in the anterior pretectal nucleus (APN) that are labelled by the *KalTA4u508* transgene and function as a command system to induce hunting routines^30^. Hunting invariably commences with a convergent saccade^18^ and accordingly, we observed many ROIs in this region of pretectum, adjacent to retinal arborisation field 7 (AF7), which showed substantial convergent saccade-triggered modulation of calcium fluorescence [Fig.7A]. Because optogenetic activation of APN command neurons is sufficient to trigger the production of convergent saccades, in addition to other motor components of hunting routines, and a subset of these cells project to the mid/hindbrain^30^, we investigated whether they might make synaptic connections with any of the circuit elements involved in convergent saccade generation.

**Figure 7:**
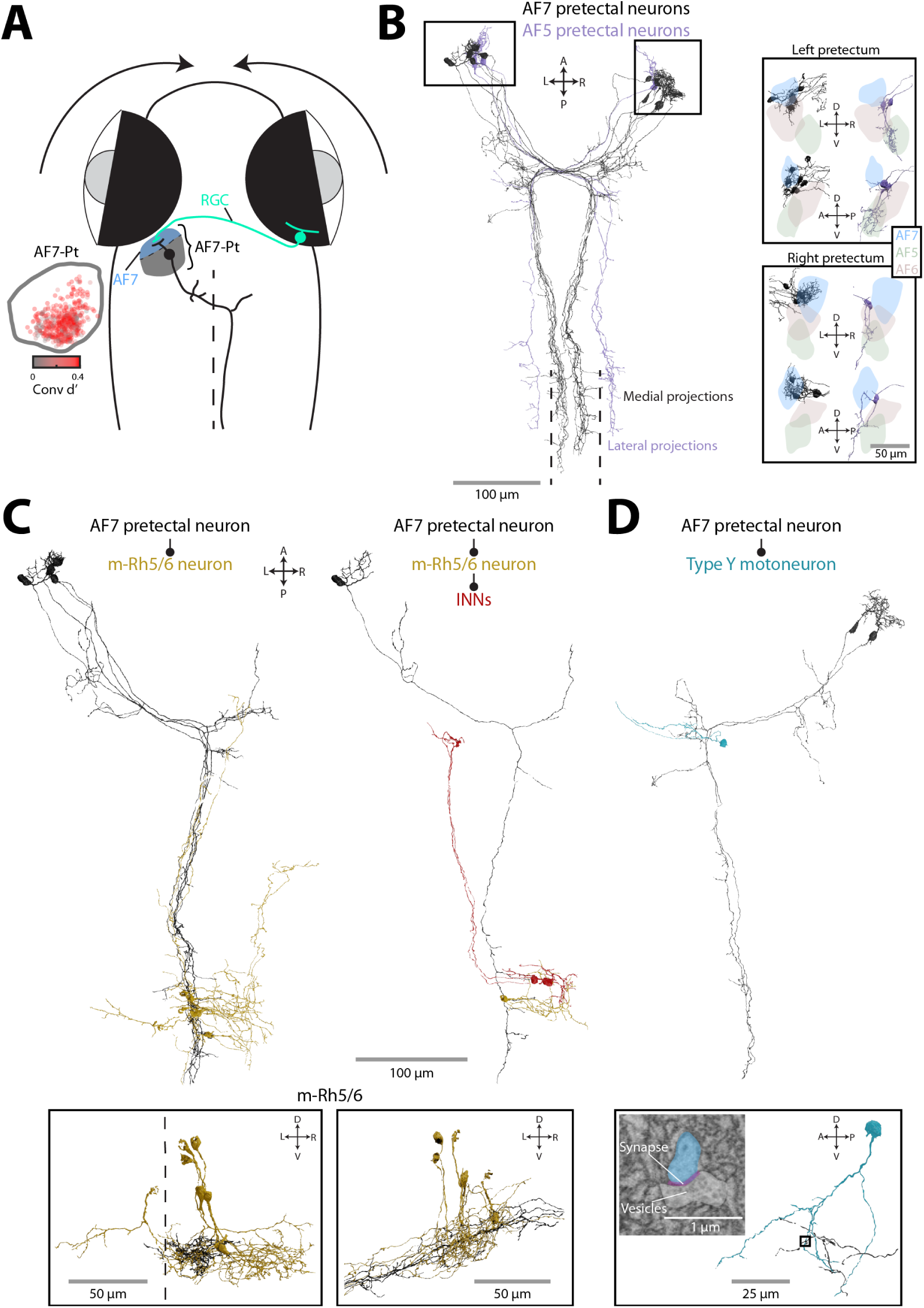
Descending commands from pretectum. (A) Schematic of retinal arborisation field 7 (AF7), adjacent pretectum (AF7-Pt) and an APN command neuron (black). *Inset:* Convergent saccade-triggered activity modulation (Conv d’) for oculomotor-tuned ROIs in AF7-Pt (485 neurons from 30 fish). (B) Ultrastructural reconstructions of pretectal projection neurons (12 AF7-pretectal cells and 3 AF5-pretectal cells). Main panel shows horizontal projection and inset boxes show coronal and sagittal views of dendritic arbors relative to retinal arborisation fields (AF5–7). (C) Horizontal projection of three AF7-pretectal neurons and six post-synaptic cells in m-Rh5/6. Inset boxes show coronal and sagittal views of m-Rh5/6. (D) Two AF7-pretectal neurons and post-synaptic Type Y motoneuron (caudal extent of the pretectal neuron axons has been cropped). Inset box shows an electron micrograph and location of one of the synapses.

Indeed, ultrastructural reconstructions revealed that AF7-pretectal neurons made synaptic connections with targets in m-Rh5/6 as well as with Type Y MRMNs. We examined 12 AF7-pretectal neurons from the Svara et al, 2022^40^ EM dataset whose soma locations and axo-dendritic morphologies matched those described for *KalTA4u508* APN command neurons^30^ [Fig.7B]. These neurons extended dendrites into AF7 [Fig.7B inset] and projected long axons that decussated in the vicinity of the oculomotor nucleus and then extended caudally in the contralateral hindbrain reticular formation, very close to the midline (*∼* 5 *µ*m from the mid-line). We identified synaptic connections between the axons of these putative APN command neurons and cells in m-Rh5/6 [Fig.7C], with one post-synaptic neuron itself contacting two INNs [Fig7C right]. Furthermore, we identified synaptic connections from two AF7-pretectal neurons onto the distal dendrite of a contralateral Type Y motoneuron, indicating that descending pre-tectal commands directly impinge on the extraocular motoneurons that our data suggest are preferentially active during convergent saccades [Fig.7D].

We additionally reconstructed a second type of pretectal neuron, whose afferent and efferent connectivity suggested a role in controlling conjugate eye movements. These neurons (*N* = 3) extended dendrites into retinal arborisation field 5 (AF5), which contains the axon terminals of direction-selective retinal ganglion cells ^42^ and projected axons that decussated near nIII and coursed through the contralateral hindbrain tegmentum at a lateral position, *∼* 50 *µ*m from the midline [Fig.7B]. We identified synaptic connections from these AF5-pretectal neurons onto the somata of both INNs and LRMNs as well as neurons in dorsal m-Rh5/6 [Fig.S7]. Neurons in this region of the larval zebrafish pretectum are believed to play a similar role to the mammalian accessory optic system (DTN/NOT) and process whole-field visual motion stimuli to control the OKR and optomotor response^42–44^. To our knowledge, there is currently little evidence for direct connections from this region of pretectum to the abducens in any species ^45,46^ but the morphology and connectivity of these AF5-pretectal projection neurons appears well suited to control conjugate eye movements in response to whole-field motion.

In sum, we identified two types of pretectal projection neuron with distinct afferent inputs and efferent connectivity onto oculomotor targets. The AF7-pretectal neurons innervate m-Rh5/6 and Type Y motoneurons, compatible with commanding convergent saccades. By contrast, AF5-pretectal neurons have a connectivity pattern compatible with generation of conjugate eye movements.

### A model for saccade type-specific recruitment of MRMNs

Our findings support a model in which two types of adducting saccade, with distinct kine-matics and ethological roles, are controlled by parallel premotor pathways to produce saccade type-specific recruitment of medial rectus motoneurons [Fig.8]. We propose that giant-synapse MRMNs, located in dorso-medial nIII, are active for both conjugate and convergent adducting saccades. This motoneuron subtype is recruited by contralateral INNs which in turn are inner-vated by neurons in dorsal m-Rh5/6, forming a *saccade type-agnostic pathway*. For convergent adducting saccades, a parallel *convergent saccade-specific pathway* leads to additional recruit-ment of Type Y MRMNs located in dorso-lateral nIII. Our data suggest Type Y motoneuron activity is influenced by a combination of afferent inputs including from INNs, ipsilateral ventral m-Rh5/6, and pretectal (APN) command neurons.

**Figure 8:**
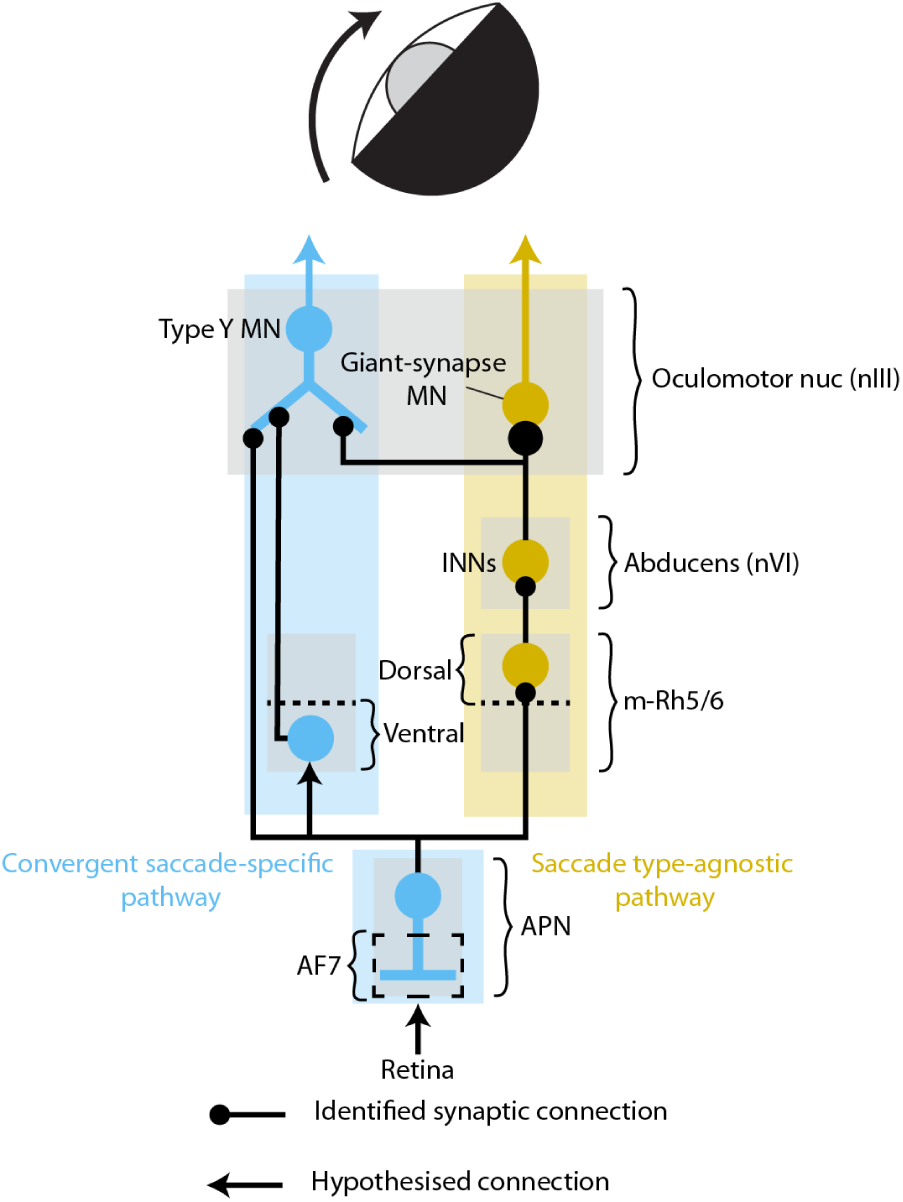
Circuit model. Two types of saccade are controlled by parallel premotor pathways that activate subsets of motoneurons: A *saccade type-agnostic pathway* (yellow) and a *convergent saccade-specific pathway* (blue). Both conjugate and convergent adducting saccades involve activation of the type-agnostic pathway in which INNs receive input from dorsal m-Rh5/6 and in turn provide giant-synaptic input to GS MRMNs in dorso-medial oculomotor nucleus. In the convergent saccade-specific pathway, Type Y MRMNs in dorso-lateral nIII are additionally recruited, likely by a combination of INN, ventral m-Rh5/6, and pretectal innervation. Descending commands from APN command neurons activate both pathways to produce convergent saccades in the context of hunting.

## Discussion

Our findings suggest that selective activation of subsets of extraocular motoneurons generates two types of kinematically distinct saccadic eye movement and we identify parallel premotor pathways that are likely to control these rapid eye movements in a behavioural context-specific manner.

### Saccade type-specific recruitment of extraocular motoneurons

For animals to move the same body part (plant) with distinct kinematics, the nervous system must generate distinct patterns of ensemble motor unit activity. In the oculomotor system, there has been longstanding debate about the extent to which extraocular motoneurons might selectively participate in certain types of eye movement^47^. Several recording studies lent support to the idea of a ‘final common pathway’ by showing that all extraocular motoneurons participate in all types of eye movement (including saccades, slow vergence movements and fixations) ^9,48–50^. However, extraocular motoneurons collectively display a broad range of physiological properties^31,51^ even to the extent that functional subtypes have been identified^13^, suggesting a degree of selective engagement according to the kinematic requirements of a specific eye movement. The concept of a final common pathway also appears at odds with the fact that there is conspicuous hetereogeneity in the structure, biochemistry and physiology of extraocular muscle fibres and their associated motoneurons. Specifically, extraocular muscles contain both fast-twitch singly innervated fibres (SIFs) and non-twitch multiply innervated fibres (MIFs), which are innervated by SIF and MIF motoneurons respectively, with distinct neurochemistry, locations within the motor nuclei, and patterns of afferent innervation (reviewed in^10^).

In this study, we used cellular-resolution calcium imaging during naturalistic behaviour to provide direct evidence that subsets of extraocular motoneurons are selectively engaged during two types of saccadic eye movement. Their locations in the dorsal subdivision of nIII, activity during adduction of the ipsilateral eye, and positional overlap with neurons we reconstructed from EM data that receive monosynaptic INN input, collectively provides strong evidence that these cells are MRMNs^31,32,35,52,53^. Our data support the notion that motoneuron recruitment is explicitly influenced by saccade *type*, independent of kinematic differences between the saccades we sampled. Specifically, the topography of MRMN activity, wherein dorso-medial MRMNs were active for both conjugate and convergent saccades whereas dorso-lateral MRMNs are specifi-cally recruited during convergent saccades, was consistent across a range of saccade amplitudes (and peak velocities, due to the main sequence relationship). Moreover, by comparing activity for kinematically matched pairs of convergent and conjugate adducting saccades, we showed at the single-neuron level that activity is modulated as a function of saccade type.

### A synaptic mechanism for differential recruitment of medial rectus motoneurons

Ultrastructural data indicated that differences in synaptic connectivity likely contribute to saccade type-specific recruitment of extraocular motoneurons. We discovered three subtypes of MRMN, with distinct dendritic morphologies and patterns of synaptic connectivity with INNs and which occupied different positions in dorsal nIII, aligned with the functional topography revealed by calcium imaging. Most surprising was the discovery of the giant-synapse motoneurons, which were located in dorso-medial nIII and whose somata enveloped a single, enormous presynaptic bouton, in a one-to-one connectivity pattern with a presynaptic abducens internuclear neuron. To our knowledge, this remarkable synapse has not previously been described in any species. The massive presynaptic terminal, containing abundant vesicles, mitochondria and multiple active zones, is reminiscent of the Calyx of Held in the auditory brainstem^54^ and is likely adapted to provide a fast and reliable feedforward relay of INN activity. This is compatible with the fact that INNs show similar activity to extraocular motoneurons, including pulse-step signals during saccades^55^ and agrees with the similarity in functional PC1 scores between dorso-medial MRMNs and INNs. Type X and Type Y motoneurons also received synaptic input from INNs, but had much larger and more complex dendritic arbors, suggesting their recruitment is influenced by other afferent pathways. Indeed, for Type Y motoneurons, we observed several additional sources of innervation, including from ventral m-Rh5/6 and pretectum. Based on these results, we propose that specific patterns of afferent innervation underlie saccade type-specific recruitment of MRMNs.

In primates, tracing studies have identified three pools of MRMNs in and around nIII^56^ that differ in size, afferent innervation and synapse organisation, leading to the suggestion that they might play distinct oculomotor roles^37,57^. MIF motoneurons reside specifically in the ‘C-group’ and it has been suggested that these cells are specialised for slow vergence eye movements^58^. Due to limited size of the serial-blockface EM volume^40^, we were unable to fully reconstruct MRMN axons and their terminations on muscle fibres and so we could not identify the *en grappe* or *en plaque* synapses that are characteristic of MIF and SIF innervation, respectively ^59^. Future studies will establish which of the three subtypes of zebrafish MRMN correspond to MIF and SIF motoneurons and how saccade type-specific neural activity and kinematics relate to differences in activation of distinct muscle fibre types. In any case, our data suggest the subdivision of MRMNs into three subtypes with specialised functions might be widely conserved across modern vertebrates, from fish to primates.

### Pathway activity and implications for movement kinematics

How might activity in the saccade type-agnostic and convergent saccade-specific pathways relate to the kinematics of adducting saccades that follow distinct main sequence relationships? If convergent saccades of all sizes involve the additional recruitment of Type Y MRMNs, it might seem necessary that during small convergent saccades there would be reduced pulse activity in the type-agnostic pathway, to account for their reduced velocity as compared to conjugate saccades of equivalent amplitude. This is certainly plausible. While we named the pathway ‘type-agnostic’, the signals carried by premotor and motor neurons might nonetheless vary across saccade types (thus representing a hybrid of the two models in [Fig.1A]). Indeed, we observed negative saccade type indices for some cells in dorso-medial nIII, nVI and dorsal m-Rh5/6, indicating elevated activity during conjugate saccades. However, it should be noted that extraocular muscle force may not easily be inferred from changes in motoneuron activity^12,60^. MIFs and SIFs have distinct force generation profiles and orbital MIFs act on pulleys to change the pulling direction of extraocular muscles such that coactivation of MIFs and SIFs might translate in complex ways to ocular kinematics^16,61^. Furthermore, recordings from MRMNs in cats revealed differences in the intra-saccadic timing of pulse activity on MIF versus SIF motoneurons, suggesting they differentially contribute to saccade kinematics^31^. In this study, the limited resolution of calcium imaging prevented us from evaluating pulse-step activity and future work will be needed to establish the firing profiles of giant-synapse and Type Y MRMNs, ideally alongside measurement of muscle force.

### Premotor control of adducting saccades in different behavioural contexts

In our model, both types of adducting saccade depend upon giant-synapse MRMNs being recruited by contralateral INNs. INNs have been long understood to mediate conjugate eye movements by coupling activity in the abducens to contralateral oculomotor nucleus to produce coordinated ipsiversive rotation of both eyes ^35,55^. While physiological recordings and lesion studies indicate that INNs are not required for slow vergence movements in mammals^61–63^, we show that they are essential for fast convergent saccades in zebrafish. This result necessitates that activity of INNs be uncoupled from LRMNs and indeed, our calcium imaging revealed INN recruitment during both saccade types, but minimal activity in LRMNs during convergent saccades when the ipsilateral eye must rotate nasally. This independence of the two neuronal types in abducens is supported by a recent connectomics study, which identified two oculomotor submodules in the zebrafish tegmentum that are preferentially connected to either INNs or LRMNs^64^. Furthermore, several studies in mammals and fish have described monocular encoding in various oculomotor cell types, including the excitatory burst neurons (EBNs) that encode saccade velocity^28,53,65–69^. In zebrafish, a likely candidate for these EBNs are the neurons we reconstructed in dorsal m-Rh5/6. Optogenetic mapping in larval zebrafish has previously identified rhombomere 5 as a locus capable of eliciting horizontal saccades^27,41^ and like EBNs in mammals^70–72^, dorsal m-Rh5/6 neurons are located in the pontine reticular formation (PRF) medial to nVI and make monosynaptic connections onto INNs and LRMNs. In line with our model, precise multiphoton optogenetic stimulation of small groups of cells within m-Rh5/6 reliably evoked adduction of the contralateral eye. Moreover, different stimulation sites showed substantial variability in their effects on the ipsi-versus contralateral eye, in line with monocular encoding in EBNs^65^. Although optogenetically evoked eye movements were slower than saccades, electrical stimulation of PRF in mammals also tends to produce rather slow, constant velocity eye rotations, which has been interpreted to be a consequence of peripheral oculomotor circuits mediating a high degree of activity integration^73,74^. Indeed, strong interconnectivity of this region with the horizontal velocity-to-position neural integrator (hVPNI) is suggested by both physiological and connectomics data in zebrafish, which indicate that highly recurrent hVPNI circuits extend through the tegmentum from rh7-8 as far rostrally as rh4-6^64,75–78^.

Zebrafish generate conjugate saccades in several behavioural contexts including to visually scan their environment when stationary, shift or maintain gaze during locomotion, and recentre the eye during OKR^17^. All conjugate saccades conform to the same main sequence relationship and so it is likely that saccadic commands from several brain regions converge upon dorsal m-Rh5/6 EBNs. Here, we identified AF5-pretectal projection neurons, which synapsed onto m-Rh5/6 neurons as well as LRMNs and INNs. It is unclear if these cells command saccades or other types of conjugate eye movement. Given that optogenetic stimulation of this region of pretectum (and electrical stimulation of its supposed mammalian equivalent) evokes slow phase eye movements^43,79^, it is possible that AF5-pretectal neurons might mediate the early direct component of the OKR in which the eye responds to the onset of visual motion with a rapid increase in slow phase velocity^80,81^. A recent study discovered a population of hindbrain neurons that display pre-saccadic ramping activity, which predicts the occurrence of spontaneous saccades^29^. In future work, it will interesting to determine if and how these cells, as well as other afferent neurons, interface with the saccade type-agnostic pathway we have proposed.

Convergent saccades are used by zebrafish to binocularly foveate their prey and switch into a predatory mode of gaze during hunting^17,18^. We propose that this saccade type is generated by additional activation of a parallel premotor pathway that recruits Type Y MRMNs. Previously, we showed that APN neurons function as a hunting command system^30^ and here we show that they directly synapse onto Type Y MRMNs, providing a link between pretectal induction of hunting state and hunting-specific oculomotor output. Although the signals carried by these cells are not yet known, the fact that their optogenetic activation is sufficient to induce convergent saccades, along with their connectivity to m-Rh5/6, suggests they might function as long-lead burst neurons, providing phasic activity that recruits EBNs. Type Y cells also received an ipsilateral input from neurons in ventral m-Rh5/6, which were tuned to adduction of the ipsilateral eye and showed convergence-specific activity. At this stage we can only speculate as to the function of these cells. In mammals, ipsilateral ascending inputs to MRMNs derive from ascending tract of Dieters (ATD) neurons in lateral and medial vestibular nucleus (LV, MV) ^34^ and the nucleus prepositus hypoglossi (NPH)^82^, which forms part of the hVPNI. The cells we described are located only *∼* 25 *µ*m from the midline, indicating they are unlikely to be part of MV^83^ and are clearly too medial to reside in LV. Furthermore, recordings in cat found that NPH neurons innervating nIII have an ON-direction corresponding to abduction of the ipsilateral eye, opposite to our results. Therefore, it is possible the cells we identified represent a previously undescribed input to MRMNs. Their location in m-Rh5/6 suggests they may function as burst neurons, providing Type Y cells with an eye velocity signal, while the putative relationship with NPH raises the possibility that they (perhaps additionally) carry an eye position signal. Zebrafish sustain high ocular vergence for the duration of hunting sequences^18^. The hVPNI (or possibly a separate vergence integrator) is expected to generate the necessary eye position signals and these might reach Type Y neurons via this pathway from ventral m-Rh5/6. We note that sustained activity in APN neurons could also contribute to maintaining or adjusting post-saccadic eye position. Future work will clarify the signals carried by these premotor elements and resolve how they cooperate with INN innervation to recruit Type Y MRMNs during predatory eye convergence.

### Brainstem control of fast vergence

How does our model compare to what is understood of fast vergence control in primates? The convergent saccades of larval zebrafish^17^ are similar to disjunctive saccades (DS) of primates^84^, which also comprise version (conjugate) and vergence components to shift fixation between targets in 3D space. Although these are by far the most common saccades used in everyday viewing, their neural basis has received little attention as compared to conjugate saccades and slow, symmetric vergence (driven by accomodative and disparity signals) ^85^. Previous models have sought to explain DS by combining activity of the (conjugate) saccade and vergence subsystems in a schema that accords with Hering’s ideas about binocular coordination of eye movements, in which identical commands are sent to both eyes ^86^. To account for the high vergence velocities that are obtained during DS, such models hypothesised the existence of an additional neural component, saccade-vergence burst neurons (SVBNs), and recently cells carrying the appropriate signal were discovered in the central mesencephalic reticular formation of monkey^87^. We were not able to identify cells with activity similar to SVBNs, nor cells encoding vergence position or velocity^88–91^, suggesting zebrafish are unlikely to have a midbrain vergence subsystem. However, recent work has challenged the idea that such a system is required for DS, perhaps even in primates. Specifically, the finding that a wide variety of neurons in the ‘conjugate’ saccadic pathway appear to carry monocular signals supports a model in which each eye is programmed independently and as such the saccadic system can mediate both the version and vergence components of DS ^85^. Our data support and extend this model by proposing that both the canonical saccadic pathway, as well as an additional parallel pathway, act together to generate fast vergence eye movements. The circuits we have mapped in larval zebrafish might represent an ancestral vertebrate blueprint for the control of fast vergence and later in evolution, the appearance of a midbrain vergence system may have arisen for more precise binocular foveation and stereopsis^61^. Finally, although we define convergent and conjugate saccades in a binocular context, here we have outlined pathways that control adduction of a single eye. In future work, we hope to leverage the experimental accessibility of the larval zebrafish brain to understand how the animal coordinates both eyes to binocularly foveate its prey.

## Acknowledgements

The authors thank members of our lab, Troy Margrie, Josh Bassett and Vanessa Ruta for helpful discussions and feedback on the project, UCL Fish Facility staff for fish care and husbandry, and Fabian Svara and Dominique Foerster for help using the EM dataset. This research was funded in whole, or in part, by the Wellcome Trust (Grant numbers 101195/Z/13/Z and 220273/Z/20/Z awarded to I.H.B.). For the purpose of Open Access, the author has applied a CC BY public copyright licence to any Author Accepted Manuscript version arising from this submission.

C.K.D. was supported by a Wellcome Trust 4 year Neuroscience PhD studentship.

## Author Contributions

Conceptualisation and Methodology: C.K.D. and I.H.B.; Investigation: C.K.D., T.H.; Analysis: C.K.D.; Writing: C.K.D. and I.H.B.; Supervision and Funding Acquisition: I.H.B.

## Declaration of Interests

The authors declare no competing interests.

## Methods

### Animals

Zebrafish lines were maintained in the Tübingen background. Larvae were reared in fish-facility water on a 14/10 h light/dark cycle at 28.5*^◦^*C and were fed *Paramecia* from 4 dpf onwards. All larvae were homozygous for the *mitfa^w^*^2 92^ skin-pigmentation mutation. For functional imaging, animals were transgenic for Tg(*elavl3*:H2B-GCaMP6s)jf5Tg ^93^. Photoactivations were performed using larvae carrying Tg(*Cau.Tuba1*:c3paGFP)a7437Tg ^94^ and Tg(*elavl3*:jRCaMP1a)jf16Tg ^95^. Optogenetic stimulation was conducted using Tg(KalTA4u523);Tg(UAS:CoChR-tdTomato)u332Tg (below and^96^). The sex of the larvae is not defined at the early stages of development used for these studies. Experimental procedures were approved by the UCL Animal Welfare Ethical Review Body and the UK Home Office under the Animals (Scientific Procedures) Act 1986.

### Generation of transgenic zebrafish

The Tg(–2.5*pvalb6*:KalTA4)u523Tg [abbreviated Tg(KalTA4u523)] transgenic line was isolated by screening the progeny of animals injected with a –2.5*pvalb6*:KalTA4 expression construct that was generated and injected as described in^30^. This expression vector generated a wide range of expression patterns, one of which was designated the allele u523Tg and labelled a broad population of neurons in midbrain and hindbrain.

### Two-photon calcium imaging and behavioural tracking

Larvae were tethered in 3% low-melting point agarose gel in a 35 mm petri dish lid and sections of gel were carefully removed using an opthalmic scalpel to allow free movement of the eyes and tail below the swim bladder. Larvae were allowed to recover overnight before testing at 6 or 7 dpf. Imaging was performed using a custom-built multiphoton microscope [Olympus XLUMPLFLN *×*20 1.0 NA objective, 580 nm PMT dichroic, bandpass filters: 510/84 (green), 641/75 (red) (Semrock), R10699 PMT (Hamamatsu), Chameleon II ultrafast laser (Coherent)] at 920 nm with laser power at sample of 5–10 mW. Images (0.67 *µ*m/px) were acquired by frame scanning at 4.8 Hz, with focal planes separated by 10*µ*m.

Eye position was monitored at either 60 or 300 Hz using a FL3-U3-13Y3M-C camera (Point Grey) that imaged through the microscope objective under 720 nm illumination. Tail position was imaged at 420 Hz under 850 nm illumination using a sub-stage GS3-U3-41C6NIR-C camera (Point Grey). Horizontal eye position and tail posture (defined by 13 equidistant x-y coordinates along the anterior-posterior axis) were extracted online using machine vision algorithms^97^.

Two projectors were used to present visual stimuli. The first (Optoma ML750ST) back-projected stimuli onto a curved screen placed in front of the animal at a viewing distance of 35 mm while the second (AAXA P2 Jr) projected images onto a diffusive screen directly beneath the chamber. Visual stimuli were defined using the ‘red’ colour channel and Wratten filters (Kodak, no. 29) were placed in front of both projectors to block residual light that might be detected by the PMT. Visual stimuli were designed in MATLAB using Psychophysics Toolbox^98^. Prey-like moving spots comprised 5*^◦^* bright or dark spots (Weber contrast +1 or -1 respectively) moving at 30*^◦^*/s either left*→*right or right*→*left across 152*^◦^* of frontal visual space. Optokinetic stimuli were presented in front of the animal and comprised drifting sinusoidal gratings (wavelength 19*^◦^*, velocity 0.3 cycles/s, Michelson contrast 0.5) that alternated between leftwards or rightwards motion every few seconds. Stimuli were presented in a pseudo-random sequence with a 30 s inter-stimulus interval.

Microscope control, stimulus presentation and behaviour tracking were implemented using Lab-View (National Instruments) and MATLAB (MathWorks).

### Saccade detection and classification

Saccadic eye movements were analysed as per^17^. Raw eye position traces were first interpolated onto a 100 Hz time-base and low-pass filtered with a cut-off frequency of 1 Hz. Rapid eye movement events were detected as peaks in the convolution of filtered eye position with a step function (width 160 ms), with the time of the peak providing a first coarse estimate of saccade time. Rapid eye movement events of the left and right eye that occurred within 100 ms of one another were paired and treated as a single binocular event. After this pairing step, events that occurred within 300 ms of a preceding event were discarded, to limit overlap between windows for calculating saccade metrics (see below) and because manual inspection revealed these movements were rarely saccadic.

To reliably estimate eye position and velocity metrics, raw eye position traces were interpolated onto a 500 Hz timebase and smoothed with a custom LOWESS function, which was designed to reduce noise without ‘flattening’ changes in eye position during saccades. Specifically, eye position data was smoothed using the MATLAB lowess function (with span 80 ms) except for periods where the convolution of raw eye position data with a step function exceeded 3*^◦^*(putative saccades). A refined estimate of saccade onset time was then determined by convolving smoothed eye position with two step functions of width 100 ms and 40 ms, taking the product between both convolutions and thresholding the output within a 400 ms window spanning the initial estimate of saccade time.

For each rapid eye movement event we evaluated: (a) pre-saccadic eye position, as median eye position during a 200 ms window immediately prior to onset time; (b) max post-saccadic eye position, as the eye position within a 200 ms window starting at onset time that had the greatest absolute deviation from eye position at onset time; (c) median post-saccadic eye position, as median eye position over a 200 ms window starting at the timepoint corresponding to max post-saccadic eye position; (d) eye velocity (cw and ccw), as the maxima and minima, respectively, of the time derivative of eye position, determined by the MATLAB gradient function over a 150 ms window centred at onset time. We then used these measures to calculate nine oculomotor metrics describing each (binocular) rapid eye movement event: *Amplitude* (left and right eye), was the difference between median post-saccadic eye position and pre-saccadic eye position. *Max-median amplitude* (left and right eye), was the difference between max post-saccadic eye position and median post-saccadic eye position and quantifies the degree to which eye position is maintained following a saccade. *Velocity* (cw and ccw for both left and right eye), as described above. *Vergence*, was the difference between median post-saccadic eye position of the right and left eye. These metrics were normalized for each animal by winsorizing the data between the 0.5th and 99.5th percentile and then z-scoring.

To classify rapid eye movement events and label specific saccade types, we first used a MATLAB implementation^99^ of UMAP^100^ (run_umap, metric=Euclidean, min_dist=0.11, n_neighbours=199) to perform a supervised embedding into a two-dimensional UMAP space previously derived from 152 tethered animals^17^. Then, for each rapid eye movement event, a saccade-type identify was chosen by taking the modal identity of 100 nearest neighbours in the embedding space. In this study, we restricted our analysis to left conjugate, right conjugate and convergent saccades. For regression modelling of calcium time series (below), left and right lateralised convergent saccades were distinguished by the sign of post-saccadic version (the average of left and right eye position).

### Calcium imaging analysis

#### Image processing and time-series extraction

Motion correction of fluorescence imaging data was performed as per^97^. Regions of interest (ROIs), corresponding to GCaMP labelled nuclei of individual neurons, were segmented using an algorithm from^101^. The fluorescence time series of each cell was initially computed as the mean value of pixels belonging to the corresponding binary mask for each imaging frame, assigned to a time point corresponding the midpoint of the frame. For frames where motion error exceeded 5 *µ*m, pixel values were replaced by interpolation. This initial time series estimate was then detrended to correct for slow variations in fluorescence and standardised by (1) subtracting baseline fluorescence, estimated as the 50th percentile of pixel values and (2) dividing by the ‘noise’ of the calcium signal baseline, estimated using the OASIS^102^ subfunction estimate_baseline_noise. The resulting fluorescence time-series is denoted *zF*.

#### Oculomotor-tuned ROIs

Oculomotor-tuned ROIs were identified using a two-stage analysis. First, we identified ROIs as being ‘saccade-active’. Second, we identified ‘oculomotor-tuned’ ROIs as the subset of saccade-active ROIs whose fluorescence was best explained by oculomotor variables, as opposed to other stimulus or motor variables.

To identify saccade-active ROIs, we first computed for each ROI three d’ values, one for each saccade type (i.e. convergent saccades (*Conv*) and left and right conjugate saccades (*L Conj*, *R Conj*)), where

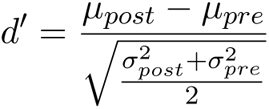

and *µ_post_* and *σ_post_*^2^ were the mean and variance of zF across time and individual instances of the saccade type during a 2 s window following saccade onset. *µ_pre_* and *σ_pre_*^2^ were computed similarly during a 1 s window prior to saccade onset. For each ROI and saccade type, a null distribution of 1000 d’ values was generated by randomising saccade onset times. An ROI was considered active for a given saccade type if the relevant d’ value exceeded the 95th percentile of this null distribution.

In the second step, we used linear regression to model *zF* for each saccade-active ROI. We designed 33 regressors, derived from 14 behavioural predictors and 18 stimulus predictors [Table S1]. Six ‘oculomotor predictors’ comprised four predictors describing the occurrence of saccades and two eye position predictors. The saccadic predictors were one-hot encodings indicating the imaging frames corresponding to saccade onset for leftward-and rightward-directed convergent and conjugate saccades. In view of the fact that many oculomotor neurons encode ipsi-or contraversive eye position, we generated rectified eye position predictors for the left and right eye, where temporal eye positions (estimated as those positions more temporal than the median position across the entire experiment) were zeroed. Locomotor predictors were one-hot encodings of the onset time of swims, specific for direction (left/right) and vigour level (1st to 4th quartile). Of the 18 stimulus predictors, two described optokinetic drifting gratings and the remainder described small moving spots. The optokinetic grating predictors were binary vectors indicating the frames during which left-or rightwards motion was presented. Small spot predictors were one-hot encodings describing a range of stimulus locations (from -60 to +60 degrees azimuth), specific for motion direction and contrast polarity. To account for calcium dynamics, regressors were generated by convolving each predictor with a calcium impulse response function (CIRF) modelled as an exponential rise and subsequent decay with time con-stants *τ_on_*= 0.2 s and *τ_off_* = 3–5 s (see below). In addition, to capture fluorescence modulations that might result from residual motion artefacts, we included a ‘motion-error’ regressor, derived from the translation applied during motion correction of image time series (this regressor was not convolved with the CIRF).

First, for each saccade-active ROI, we used ordinary least squares regression to optimise two hyperparameters: The *τ_off_*of the CIRF (3, 4 or 5 s) and a temporal offset applied to the regressors (0, 1, 2 or 3 frames). ROIs for which the best OLS model had *R*^2^ *>* 0.05 (67.8 *±* 9.3% of saccade-active ROIs, *N* = 76 fish) were then subjected to (more computationally intensive) ridge regression using the best performing pair of hyperparameters.

We used regularized linear regression with an L2 penalty (‘ridge’ regression) to model zF for selected ROIs. Ridge regression was performed using the MATLAB ridge function, with lambda selected by ten-fold cross-validation. To estimate the unique contribution of each regressor to the model fit, we followed the approach of ^103^. Specifically, for each regressor in turn, we circularly permuted the regressor by a random number of frames and recomputed the model fit using the same lambda value and cross-validation folds as per the original fit. In this way we derived a change in cross-validated goodness-of-fit (Δ*cvR*^2^) resulting from randomising the temporal relationship between the regressor in question and the recorded calcium fluorescence of the ROI. Negative values of Δ*cvR*^2^ indicate the regressor made a unique contribution to explaining ROI activity that could not be compensated by the remaining regressors. Positive values of Δ*cvR*^2^ represent random improvements in fit quality from permuting the predictor; we pooled these values across ROIs to generate a null-distribution and estimate significant values of Δ*cvR*^2^. Thus, a saccade-active ROI was classified as oculomotor-tuned if (i) the most negative Δ*cvR*^2^ value was associated with an oculomotor regressor; (ii) at least one oculomotor regressor had a Δ*cvR*^2^ more negative than the 95th percentile of the null distribution; (iii) the motion artefact predictor was *less* negative than the 95th percentile of the null. In this way we identified oculomotor-tuned ROIs whose zF time-series was best predicted by an oculomotor variable and was not explained by residual motion error. These ROIs were labelled as being active for ‘Conv’, ‘L Conj’ or ‘R Conj’ saccades, or ‘Both’ if the d’ analysis indicated significant activation for convergent as well as either left or right conjugate saccades (see Venn diagram in [Fig.S1F]).

#### Oculomotor tuning metrics

Recti-linear fits and saccade type index were determined from normalised saccade-triggered fluorescence and normalised post-saccadic eye position and eye velocity. For each ROI, normalised saccade-triggered fluorescence was calculated by first subtracting from *zF* its mean value over a 1 s window immediately prior to saccade onset and then summing *zF* over a 2 s window starting at saccade onset, thus providing a measure of saccade-triggered fluorescence change. These values were then divided by their 95th percentile across all saccades for a given ROI to provide a set of normalised saccade-triggered fluorescence values.

Post-saccadic eye position was normalised by dividing by the maximum post-saccadic nasal eye position across all saccades. Eye velocity was normalised by dividing by the 95th percentile of nasal eye velocities across saccades.

Recti-linear fits of normalised saccade-triggered fluorescence versus normalised post-saccadic eye position consisted of a horizontal baseline, equal to median fluorescence across a given span and a linear ramp, which was computed by least-squares regression starting at a threshold post-saccadic eye position. Successive fits were computed with baselines spanning progressively larger portions of the data and the fit with the lowest mean-squared error was selected if the ramping portion increased above/below threshold. Otherwise only a baseline was fit. Separate fits were made for abducting and adducting saccades for each eye.

Saccade type index was estimated by first matching conjugate saccades with kinematically similar convergent saccades for a given eye. Specifically, for each conjugate adducting saccade, a matched convergent adducting saccade was found when its Euclidean distance was *<* 0.1 in normalised post-saccadic eye position and velocity space. If more than one convergent saccade fell within this radius, the closest was selected. Next, for each ROI, we computed the median difference between normalised saccade-triggered fluorescence across these matched saccade pairs. For a proportion of ROIs (18%), saccade type index could not be computed because there were no matched pairs of saccades.

For computing saccade type indices, the activity of each ROI had to be considered with respect to either the left or right eye. To select the appropriate eye, a directionality preference was established by summing Δ*cvR*^2^ values corresponding to leftwards (LConj, ConvGL, and right eye nasal position) or rightwards (RConj, ConvGR, and left eye nasal position) eye movements; the more negative sum specified directionality preference. ROIs with a preference for leftward movement had saccade type index and nasal ramp fits computed with respect to the right eye and temporal ramp fits computed with respect to the left eye. For ROIs with rightwards preference, the eyes were reversed. In support of this approach, the best ramp fit corresponded to this directionality preference for the vast majority (83%) of ROIs.

The OKR power metric was designed to quantify activity modulation associated with slow phase eye movements during the optokinetic response. When averaged over multiple OKR presentations, eye position traces showed periodic movement that followed the direction of whole-field motion with the effects of reset saccades largely smoothed out. Thus, for each ROI, we computed the median of *zF* across presentations of leftwards and rightwards OKR gratings. Because these slow phase movements were offset by *∼* 180*^◦^*, we computed the difference between these median responses to isolate direction-selective phasic signals. We then computed the Fourier transform and assessed power spectral density at the frequency of OKR direction modulation to yield an OKR power score.

### Photoactivation of paGFP

Larvae homozygous for Tg(*Cau.Tuba1*:c3paGFP)a7437Tg ^94^ and Tg(*elavl3*:jRCaMP1a)jf16Tg ^95^ were mounted in 1% low-melting point agarose at 5 dpf. The same custom 2-photon microscope used for functional calcium imaging was used for photoactivations. paGFP was photoactivated by continuously scanning a small volume at 790 nm, 5-10 mW power at sample for 3–8 min per plane. Photoactivation volumes were 40 *×* 40 *×* 20 *−* 30 *µ*m x-y-z with focal planes spaced 5 *µ*m apart. Animals were then unmounted and allowed to recover for 24 h after which paGFP and jRCaMP1a were imaged at 1040 nm.

Locations of retrogradely labelled somata were manually determined from image volumes registered to ZBB coordinate space. The axon terminals of abducens internuclear neurons were measured using ImageJ from high-resolution stacks (0.1 *×* 0.1 *×* 1 *−* 2 *µ*m/px) in oculomotor nucleus. The sizes of axonal boutons were measured in the focal planes within which they had the largest cross-sectional area.

### Analysis of electron microscopy data

For ultrastructural reconstruction of neurons we used a publicly available serial-blockface electron micrograph volume from a 5 dpf larval zebrafish acquired at 14 *×* 14 *×* 25 nm^40^. The dataset consisted of automatically detected and over-segmented cell bodies and neurites, which we manually agglomerated into neuron morphologies using the Knossos open source software (https://github.com/knossos-project/knossos). Since our aim was to identify connections between functionally defined brain regions, we did not necessarily fully reconstruct every neuron; in any case, numerous artefacts in the dataset often precluded this.

Reconstructions of abducens internuclear neurons began either from cell bodies in the abducens nucleus, or from large axon terminals in the oculomotor nucleus. Complete axon morphologies and partial dendritic morphologies were reconstructed by merging segments in ‘Agglomeration’ mode in Knossos. Segments that terminated within a neurite and thus over-segmented the neuron, were merged. Neurites were visualised in 3 orthogonal views of the raw EM data and followed until they terminated, merged with the cell body or until we encountered an artefact in the data.

Extraocular motoneurons were reconstructed within the oculomotor nucleus (nIII) from sites post-synaptic to abducens internuclear neurons and are therefore assumed to be medial rectus motoneurons^35^. Synapses were identified according to four criteria: (1) Close apposition between the two membrane surfaces; (2) identification of punctate dark spots close to the opposing membranes, consistent with pre-synaptic vesicles; (3) darkening or thickening of the post-synaptic membrane; (4) the membrane apposition had been classified as a synapse by the automated synapse detection algorithm deployed in^40^. Reconstructed medial rectus motoneurons extended their axons into the third cranial nerve, as expected. Although we reconstructed the dendritic and axon morphologies as fully as possible, because the dataset did not encompass the extraocular muscles, it was not possible to identify post-synaptic targets or classify extraocular motoneurons as MIFs or SIFs.

Almost all neurons pre-synaptic to internuclear neurons and Type-Y motoneurons were traced from the pre-synaptic terminal (identified using the above criteria) back towards the cell body. One ventromedial rhombomere 5/6 neuron was traced from its cell body to Type Y motoneuron dendrites. Pretectal neurons were traced from the cell body as were the subset of ventral m-Rh5/6 neurons for which no connection to extraocular motoneurons was identified.

Morphologies were exported as .ply meshes and were plotted in Blender for visualisation (https://www.blender.org).

### Transmission electron microscopy

Zebrafish larvae were fixed at 6 dpf by immersion for 24 h in EM fix [2% (w/v) paraformaldehyde, 2% (w/v) EM-grade gluteraldehyde, in 0.1 M sodium cacodylate buffer (pH 7.3); all fix reagents from Agar Scientific, Stansted, UK]. Specimens were then postfixed in 1% osmium tetroxide for 3 h, dehydrated in an ethanol series, infiltrated with medium hard AGAR100 resin (Agar Scientific, Stansted, UK) and polymerised by baking at 60*^◦^*C for 24–48 h. Optimally orientated specimens were selected for sectioning from a dorsal approach. As sectioning proceeded, semi-thin (1 *µ*m thick) sections stained using Toluidine Blue were checked in batches of 5–10 until approximately the correct location was reached. This location was identified by comparing neuropil areas with the Svara et al, 2022^40^ EM dataset and ZBB light-microscopy dataset. Once the correct location was determined to have been reached, ultrathin (80 nm) sections were collected onto 2 mm Pioloform resin-coated copper slot grids (Agar Scientific, Stansted, UK). Grids were stained with uranium and lead stains and were examined using a JEOL JEM-1400Flash at 80 kV. Images were captured using a Gatan Rio16 digital camera.

### Image registration

Registration of image volumes was performed using the ANTs toolbox version 2.1.0^104^ in a similar manner to that described in^105^. Images were converted to NRRD file format for registration using ImageJ. As an example, to register the 3D image volume ‘fish1.nrrd’ to reference brain ‘ref.nrrd’, the following command was used:

antsRegistration -d 3 --float 1 -o [fish1, fish1_Warped.nii.gz] -n BSpline -r [ref.nrrd, fish1.nrrd,1] -t Rigid[0.1] -m C[ref.nrrd, fish1.nrrd,1,32, Regular,0.25] -c [200x200x200x0,1e-8,10] -f 12x8x4x2 -s 4x3x2x1 -t Affine[0.1]-m GC[ref.nrrd, fish1.nrrd,1,32, Regular,0.25] -c [200x200x200x0,1e-8,10] -f 12x8x4x2 -s 4x3x2x1 -t SyN[0.05,6,0.5] -m CC[ref.nrrd, fish1.nrrd,1,2] -c [200x200x200x200x10,1e-7,10] -f 12x8x4x2x1 -s 4x3x2x1x0

The deformation matrices computed above were then applied to any other image channel N of fish1 using:

antsApplyTransforms -d 3 -v 0 --float -n BSpline -i fish1-0N.nrrd -r ref.nrrd-o fish1-0N_Warped.nii.gz -t fish1_1Warp.nii.gz -t fish1_0GenericAffine.mat

All fluorescence imaging volumes were registered to the ZBB brain atlas^26^ and to a high-resolution reference brain [from one of the following transgenic lines: Tg(elavl3:H2B-GCaMP6s), Tg(elavl3:jRCaMP1a), Tg(elavl3:GCaMP7f) or Tg(u523:KalTA4);Tg(UAS:CoChR-tdTomato)], all with resolution 0.77*×*0.77*×*1 *µ*m/px. Registrations were conducted for different experiments as follows:

- For registration of calcium imaging data, a two-step registration process was used. First, functional calcium imaging volumes were registered to a volume of the same brain acquired at the end of the experiment with z-voxel dimension 1 *µ*m (‘post-stack’). Second, the post-stack was registered to the high-resolution Tg(elavl3:H2B-GCaMP6s) brain. Since the high-resolution brain was registered to ZBB atlas space, the transformations were con-catenated to bring the functional imaging data and ROI locations to ZBB space (calcium imaging volume *→* post-stack *→* hi-res brain *→* ZBB).
- Image volumes of retrogradely photo-labelled somata were registered to high-resolution Tg(elavl3:jRCaMP1a) or Tg(elavl3:GCaMP7f) reference brains using Tg(elavl3:jRCamp1a) expression imaged in the red channel. Transformations were applied to the green channel to bring paGFP expression into ZBB space.
- Internuclear neuron axon terminals were registered using a two-step process. A small, high-resolution volume (0.1 *×* 0.1 *×* 1 *−* 2 *µ*m/px) encompassing the axon terminals was first registered to a larger volume (308 *×* 308 *×* 50 *−* 70 *µ*m; 0.39 *×* 0.39 *×* 2 *µ*m/px) which was in turn registered to the high-resolution Tg(elavl3:jRCaMP1a) reference brain.
- For each optogenetic stimulation site a post-stimulation image was acquired. This was manually aligned to an image volume of the whole brain acquired at the end of the exper-iment, which was in turn registered to the Tg(u523:KalTA4);Tg(UAS:CoChR-tdTomato) high-resolution reference brain.

### Laser ablations

Somatic ablations were guided by anatomical location and calcium activity. Calcium imaging was performed using 6 dpf Tg(elavl3:H2B-GCaMP6s) animals and immediately after imaging maps of pixel-wise mean fluorescence modulation in response to whole-field motion, convergent and conjugate saccades were computed. Cells in the relevant anatomical region (nVI or m-Rh5/6) and which showed positive fluorescence modulations during convergent saccades, were identified. These target cells were ablated using the same custom 2-photon microscope described above, following the procedure of ^105^. In brief, animals were anaesthetised and the laser focus was spiral-scanned over the target soma for *∼* 140 ms (800 nm, 150-200 mW at sample). Ablations were deemed successful if an auto-fluorescent ‘scar’ was subsequently visible in both green and red channels. Target locations in m-Rh5/6 were logged in 10 fish (out of 13) and for nVI in 4 fish (out of 7). Following the procedure, animals were unmounted and allowed to recover overnight before undergoing subsequent functional imaging and behavioural tracking.

Axotomies were performed to sever the axons of abducens internuclear neurons in the medial longitudinal fasciculus. The axons were visualised by photoconverting paGFP in nVI at 5 dpf (see above). At 6 dpf, pre-ablation behavioural data was collected and then axotomies were performed using a similar ablation procedure, except that we used higher laser power (250-290 mW). Multiple sites were often targeted to ensure all photolabelled axons were cut. Behaviour was assayed post-ablation following overnight recovery.

### 2-photon optogenetics

Optogenetic photostimulation of CoChR-expressing neurons was performed using Tg(u523:KalTA4);Tg(UAS:CoChR-tdTomato) transgenic animals (6–7 dpf) which have widespread opsin expression in mid/hindbrain tegmentum. Stimulation was performed by raster-scanning small areas (6–20*×*6–20 *µ*m at 24–45 Hz, 920 nm, 17 mW) for 4–8 s with stimulation trials separated by 15–20 s intervals. Eye and tail movements were tracked throughout and no background illumination was provided. Changes in eye position were calculated as the difference between median eye position 250 ms prior to stimulus onset and 250 ms prior to stimulus offset. Optogenetic stimulation sites were mapped to ZBB coordinates.

### Statistical analyses

All statistical analyses were performed in MATLAB. Types of statistical test and *N* are reported in the text or figure legends. All tests were two-tailed and we report *p*-values without correction for multiple comparisons unless otherwise noted.

## Extended Data

**Figure S1:**
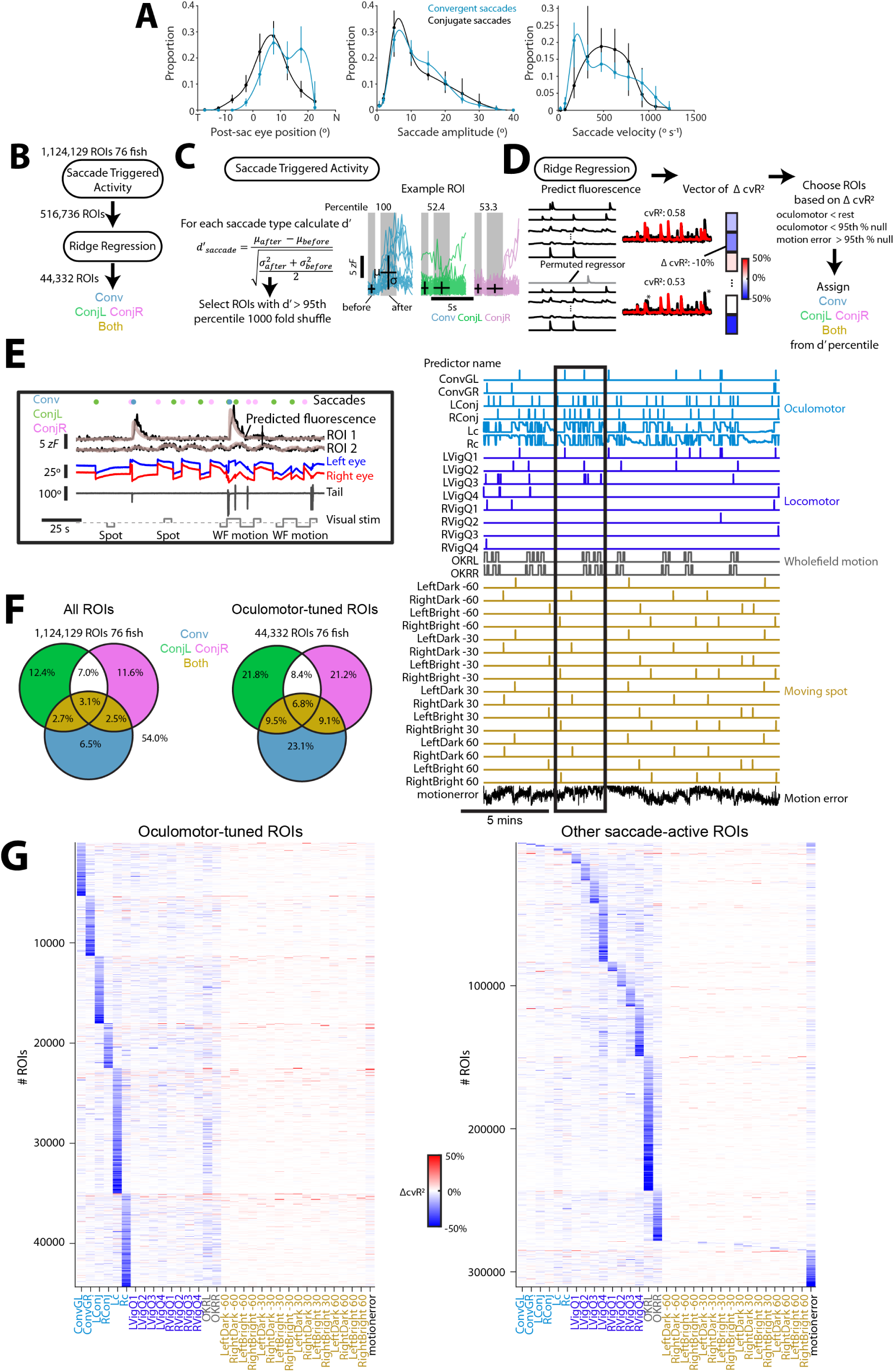
Saccade kinematics and identification of oculomotor-tuned ROIs. (A) Distributions of post-saccadic eye position, amplitude and peak velocity for convergent and conjugate adducting saccades. Median *±* IQR for *N* = 76 animals, with spline fits. (B) Overview of analysis of saccade-related activity. First, saccade-active ROIs were selected based on their saccade-triggered activity modulation (see C). Second, ridge regression was used to identify oculomotor-tuned ROIs (see D). Each oculomotor-tuned ROI was classified as *Conv*, *ConjL/R* or *Both* according to the saccade type(s) for which it was active (see F). (C) For each ROI, d’ values were computed for each saccade type and compared to null distributions calculated by shuffling saccade onset times 1000-fold. When d’ exceeded the 95th percentile of the shuffle distribution, the ROI was considered active for the corresponding saccade type. ROIs active for at least one saccade type were considered saccade-active. (D) Ridge regression was used to model the fluorescence time-series (*zF*) of each saccade-active ROI as a linear function of sensory and motor regressors. The unique contribution of each regressor to the model was quantified by circularly permuting it and assessing the fractional change in cross-validated goodness-of-fit (Δ*cvR*^2^). By repeating the process for every regressor, a vector of Δ*cvR*^2^ values is generated for each ROI. Finally, ROIs were classified as oculomotor-tuned when an oculomotor regressor produced the largest decrement in model performance (most negative Δ*cvR*^2^). For further details, see Methods. (E) Example ridge regression fits for two ROIs. The box shows recorded and model-predicted fluorescence as well as eye position, tail curvature and saccade and stimulus times during a portion of the experiment. *Right:* All 33 predictors are shown (prior to convolution with a CIRF, see Methods) for a larger portion of the experiment. Portion in the box is highlighted. (F) Venn diagrams showing the proportion of ROIs active for each saccade type, with classification key. (G) Δ*cvR*^2^ vectors for oculomotor-tuned ROIs (left) and all other saccade-active ROIs (right). ROIs have been ordered by the regressor with most negative Δ*cvR*^2^.

**Figure S2:**
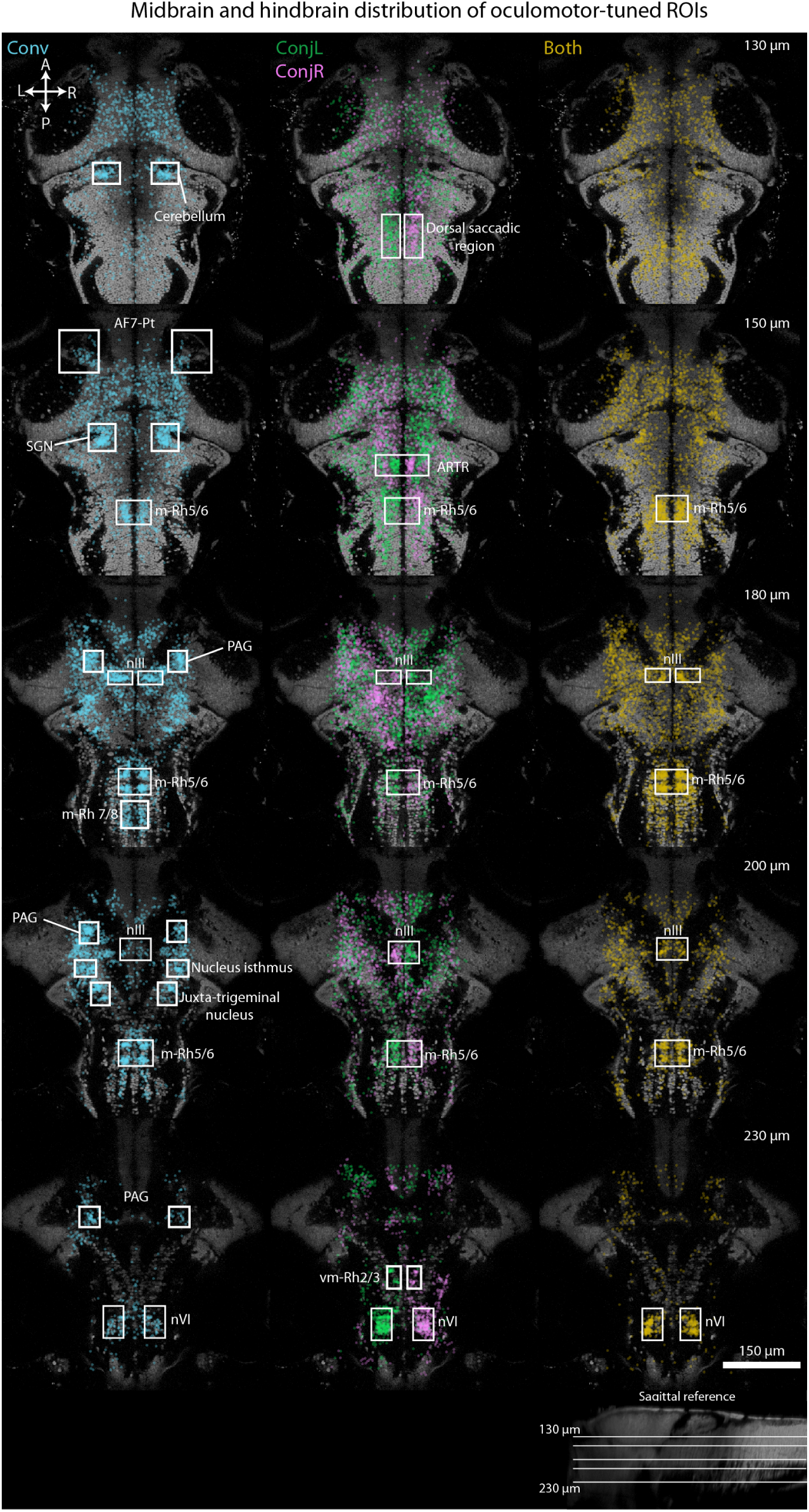
Oculomotor-tuned neurons. Oculomotor-tuned ROIs active for convergent (*Conv*) or leftwards/rightwards conjugate (*ConjL/R*) or both (*Both*) saccade types shown in ZBB reference brain space (44,332 neurons from 76 animals). Panels show horizontal planes at the dorsoventral location indicated in the top right corner and shown on a sagittal view at the bottom of the figure. All three types of oculomotor-tuned cell are found in the oculomotor and abducens nuclei as well as medial rhombomere-5/6 (m-Rh5/6), close to the facial motor nucleus. In addition, *Conj* ROIs, predominantly with ipsiversive tuning, are abundant in dorsal rhombomere 5–7, where eye velocity-related activity has been described ^27,28^, the anterior rhombencephalic turning region (ARTR) ^95^, and in ventromedial rhombomere 2/3 (vm-Rh2/3), adjacent to reticulospinal neurons. *Conv* ROIs are found in regions previously implicated in hunting, including the pretectum adjacent to retinal arborization field 7 (AF7-Pt) ^30^ and the nucleus isthmus ^105^. In addition, a high density are observed in the secondary gustatory nucleus (SGN) ^106^, the dorso-medial cerebellum, medial rhombomere 7/8 (m-Rh7/8), a tegmental region likely corresponding to the periaqueductal grey (PAG), and in close proximity to the trigeminal motor nuclei (juxta-trigeminal region). *Abbreviations*: nIII, oculomotor nucleus; nVI, abducens nucleus; ARTR, anterior rhombencepahlic turning region; SGN, secondary gustatory nucleus; vm-Rh2/3, ventro-medial rhombomere 5/6; m-Rh5/6, medial rhombomere 5/6; m-Rh7/8, medial rhombomere 7/8; PAG, periaqueductal grey.

**Figure S3:**
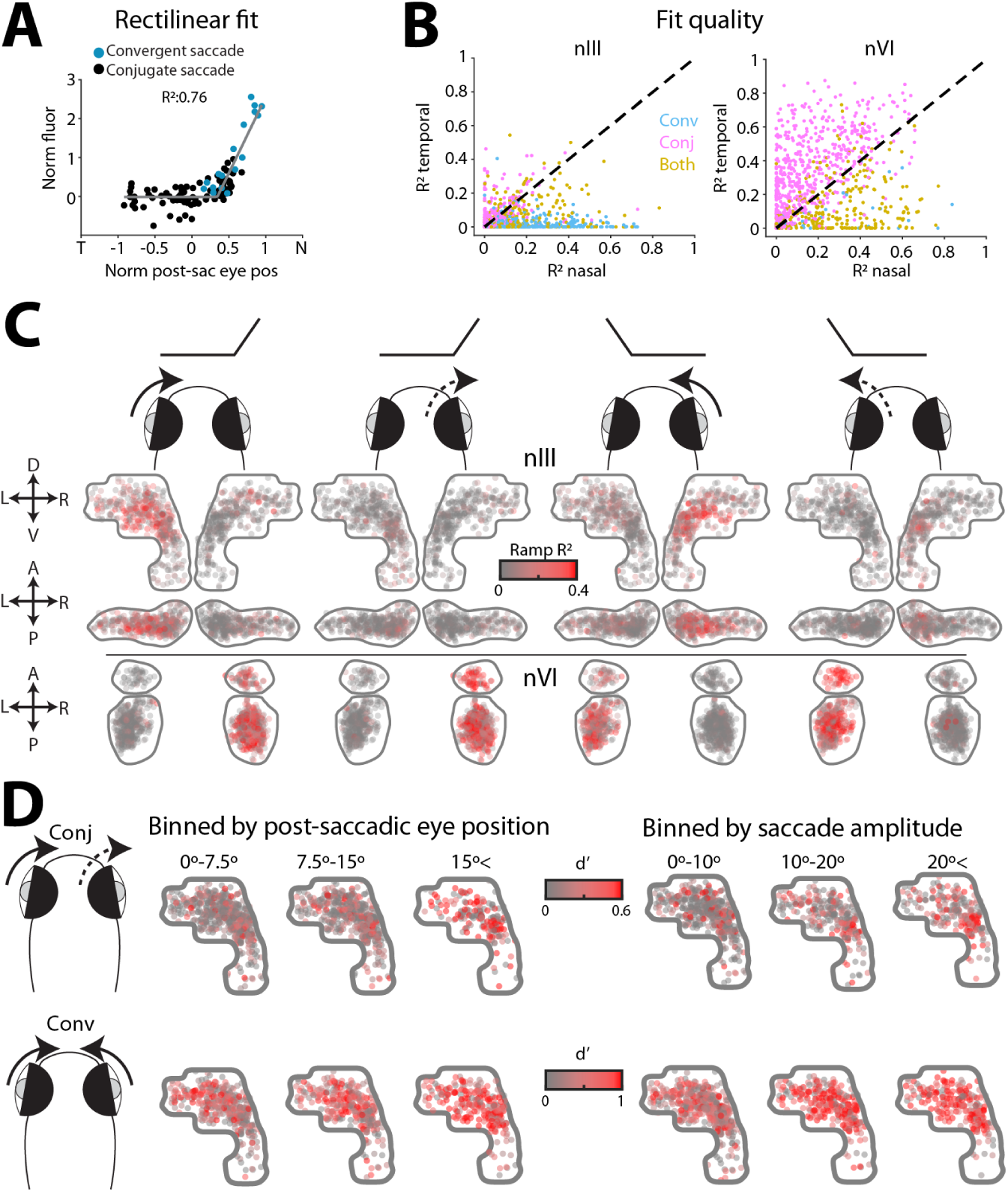
Functional properties in oculomotor and abducens nuclei. (A) Rectilinear fit of saccade-triggered change in fluorescence as a function of post-saccadic eye position, for an example neuron. (B) Comparison of *R*^2^ for rectilinear fits for nasal versus temporal eye movements in the ‘preferred direction’ of each ROI (see Methods). (C) Oculomotor-tuned ROIs colour-coded by *R*^2^ for rectilinear fits for each eye– direction contingency. From left to right: left eye nasal, right eye temporal, right eye nasal, left eye temporal. (D) Oculomotor-tuned ROIs coloured by saccade-triggered activity (d’) for conjugate and convergent saccades binned by post-saccadic eye position (left), or amplitude (right).

**Figure S4:**
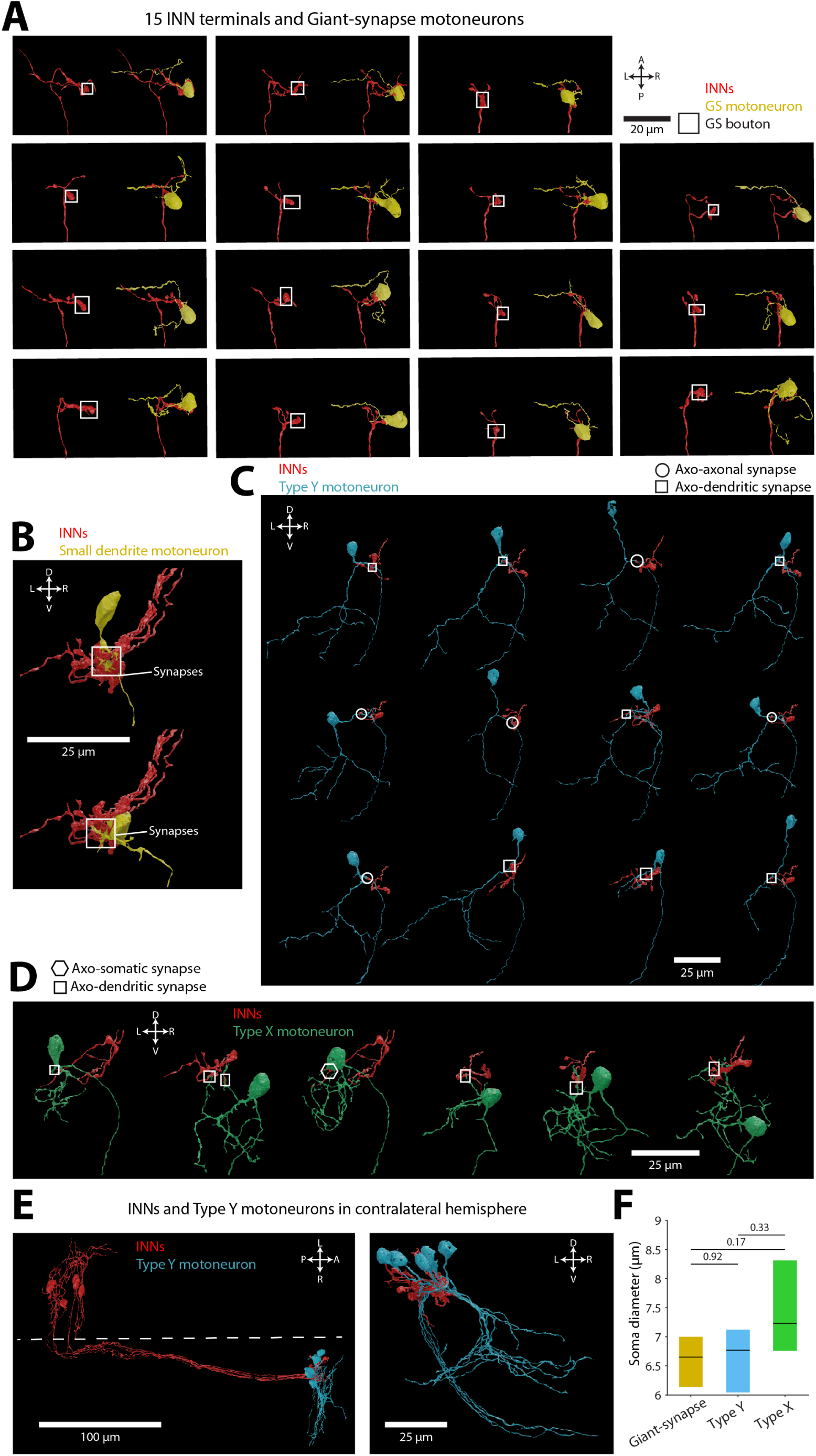
Ultrastructural reconstructions of individual motoneurons and pre-synaptic INN axon terminals. (A) 3D reconstructions of 15 giant-synapse motoneurons (yellow) and pre-synaptic INN terminals (red). Terminals forming the giant synapse indicated by white boxes. (B) 3D reconstructions of two motoneurons that formed synapses with multiple INN boutons on claw-like dendrites. (C–D) 3D reconstructions of 12 Type Y motoneurons (C) and 6 Type X motoneurons (D). (E) 3D reconstructions of an additional 6 INNs and 5 Type Y motoneurons, traced from the contralateral hemisphere. (F) Soma diameters for different motoneuron types (median (IQR) across *N* = 15 giant-synapse, 12 Type Y, 6 Type X). Kruskal-Wallis with Tukey-Kramer post-hoc tests.

**Figure S5:**
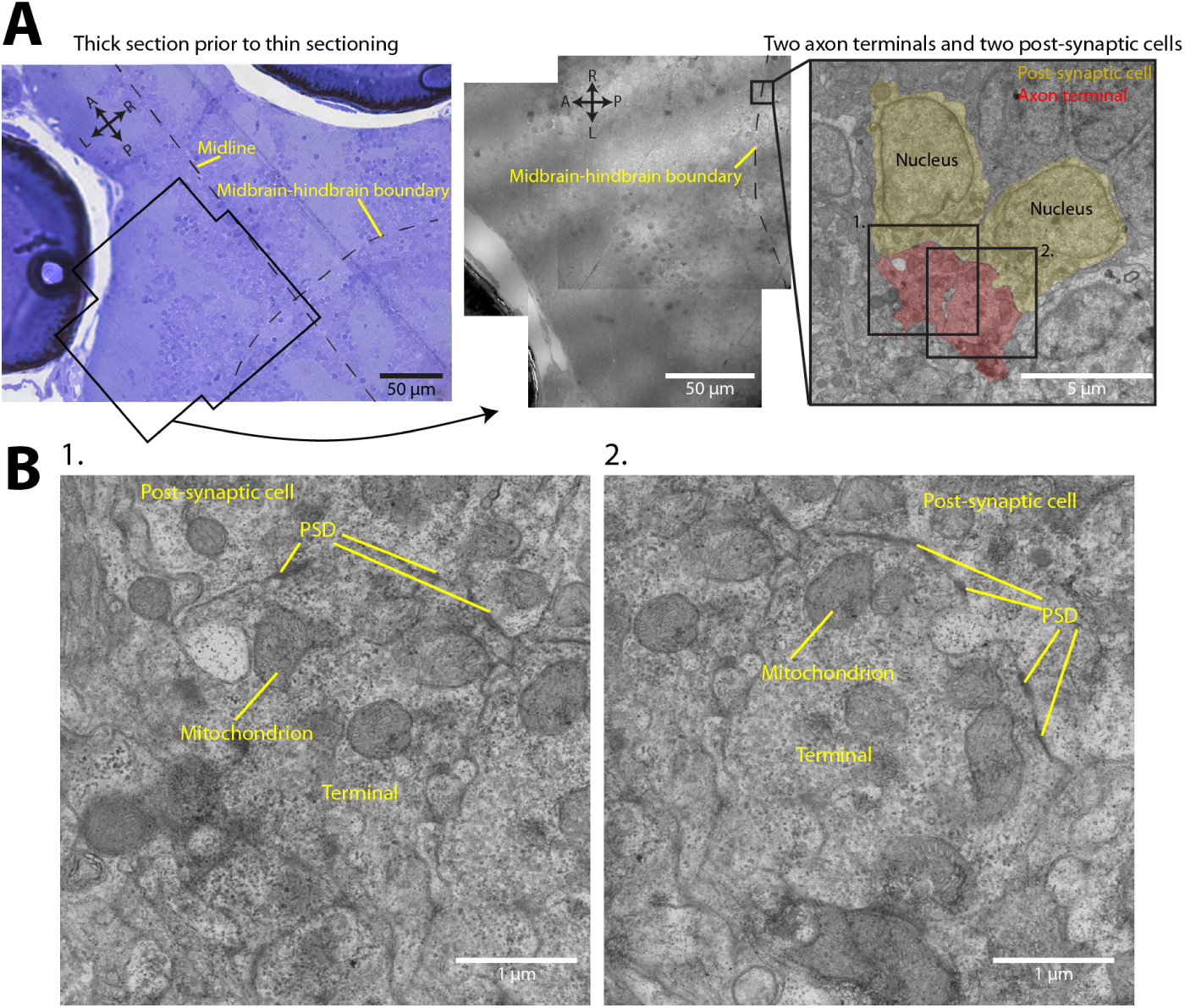
Transmission electron micrographs of giant synapses in oculomotor nucleus. (A) *Left:* Toluidine blue-stained thick (2 *µ*m) horizontal section used to guide thin sectioning for electron microscopy. Electron micrograph area shown by black outline. *Middle:* Three electron micrographs encompassing the midbrain-hindbrain boundary, aligned and overlayed. *Right:* High-magnification electron micrograph of region outlined in middle panel. Two giant axo-somatic synaptic appositions are highlighted. Boxes indicate extent of electron micrographs in B. (B) Higher magnification electron micrographs of the two giant synapses. Multiple post-synaptic densities (PSDs) can be seen at the post-synaptic membrane.

**Figure S6:**
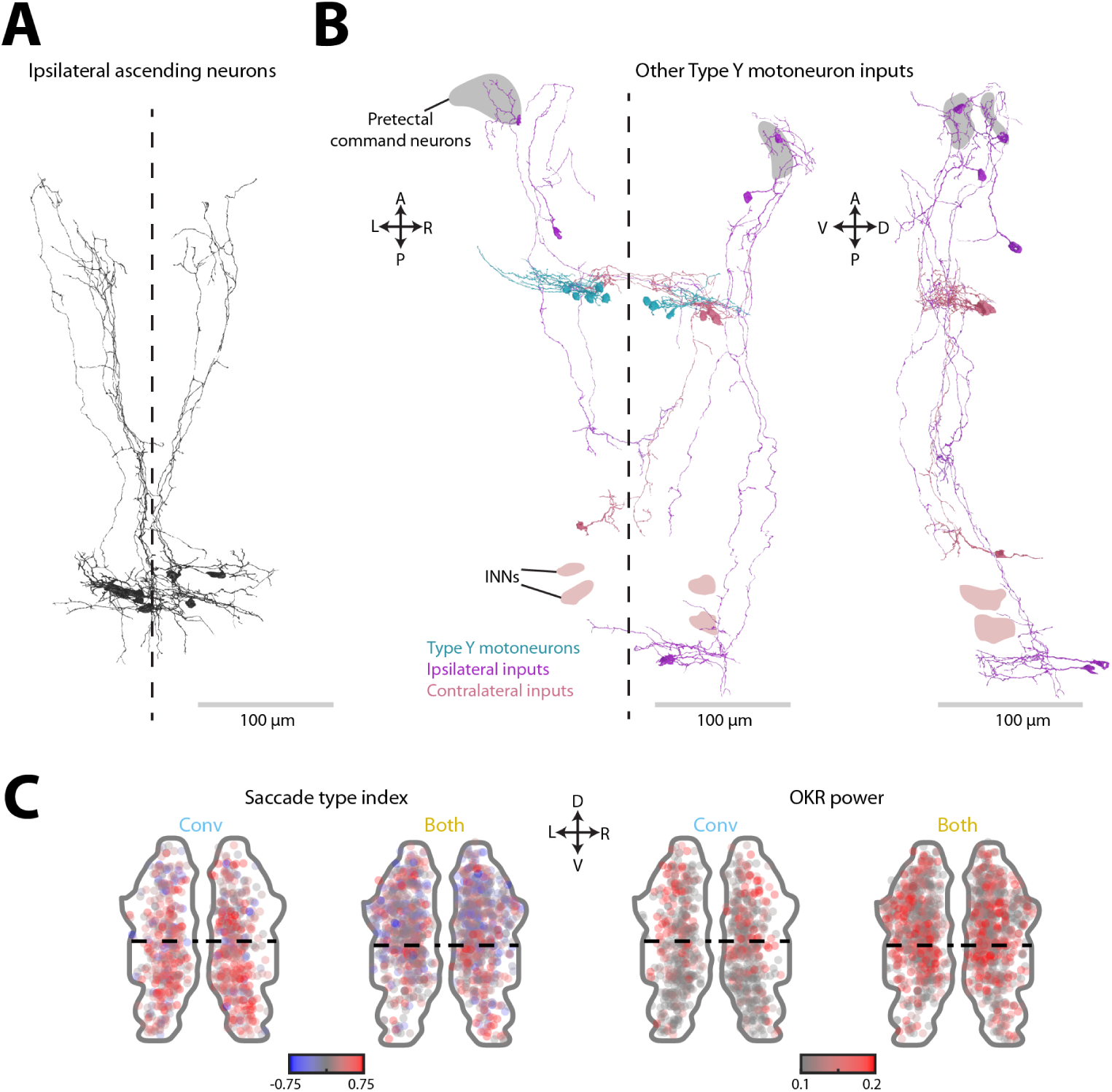
Other inputs to Type Y motoneurons and functional metrics in m-Rh5/6. (A) Ultrastructural reconstructions of nine m-Rh5/6 neurons that extended ipsilateral ascending projections to the caudal midbrain. (B) Neurons identified as presynaptic to Type Y motoneurons, other than those with somata in m-Rh5/6. Recipient Type Y motoneurons shown in the horizontal projection (left). Areas corresponding to the soma locations of INNs and pretectal command neurons are highlighted. (C) Maps of oculomotor-tuned ROIs in m-Rh5/6 colour-coded by functional metrics.

**Figure S7:**
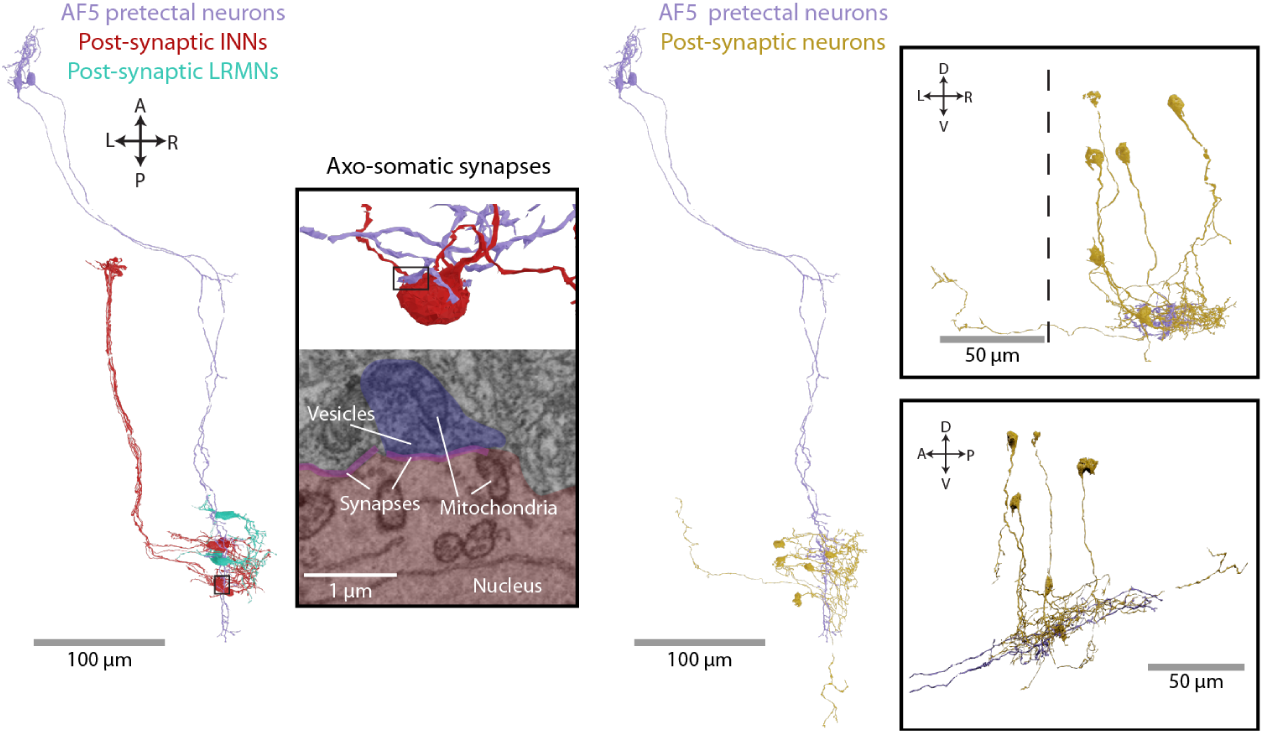
AF5-pretectal projection neurons. *Left:* Horizontal projection of two AF5-pretectal neurons with seven post-synaptic INNs and five LRMNs. Inset box shows a close-up 3D reconstruction and electron micrograph of an axo-somatic synapse onto an INN. *Right:* Horizontal projection of the same AF5-pretectal neurons along with six post-synaptic neurons in m-Rh5/6. Inset boxes show coronal (top) and sagittal (bottom) views of m-Rh5/6 region.

**Table S1:**
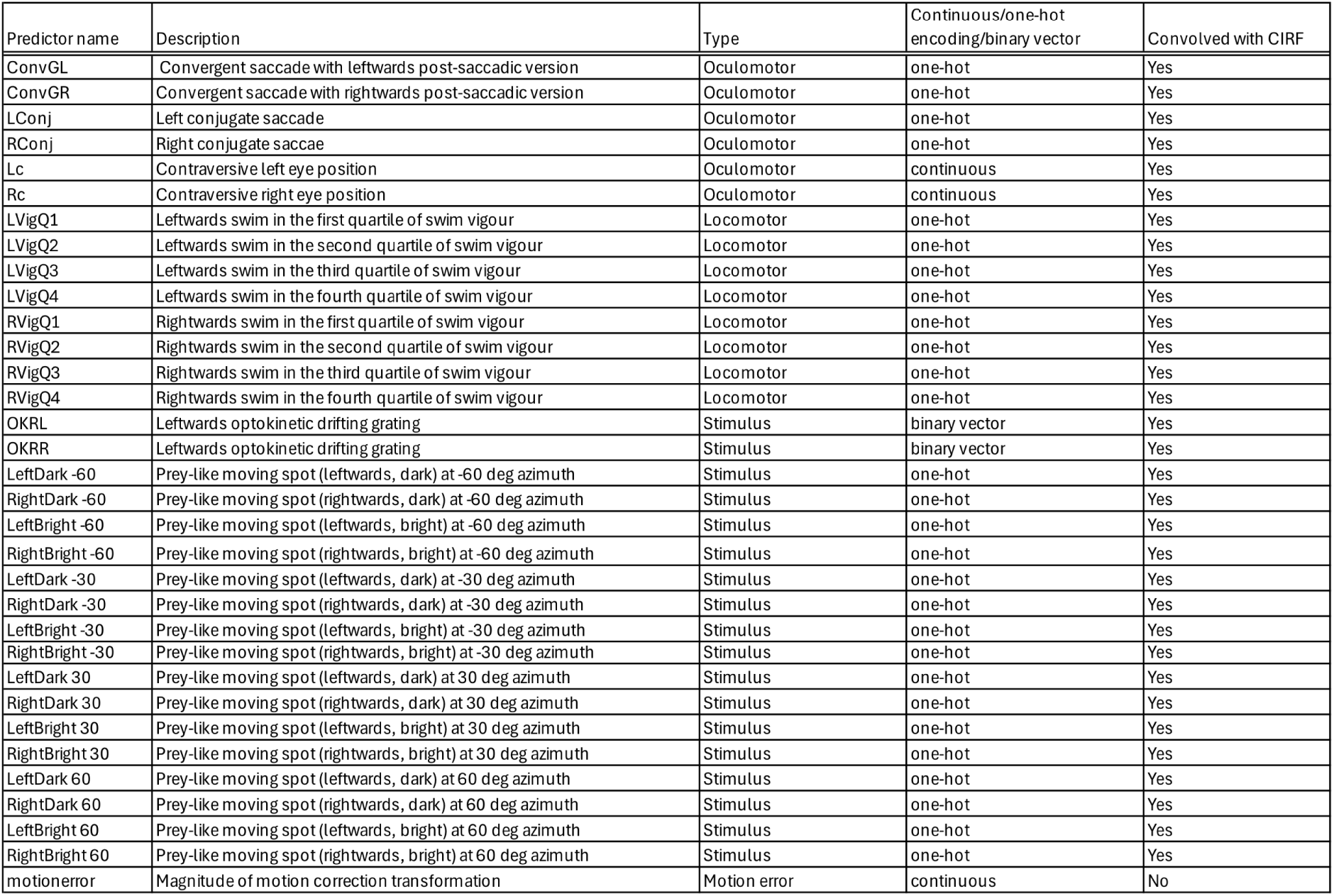
List of regressors used for linear modelling.

## Notes

### Competing Interest Statement

The authors have declared no competing interest.

